# The Power of Social Information in Ant-Colony House-Hunting: A Computational Modeling Approach

**DOI:** 10.1101/2020.10.07.328047

**Authors:** Jiajia Zhao, Nancy Lynch, Stephen C. Pratt

**Affiliations:** Massachusetts Institute of Technology, Department of Electrical Engineering and Computer Science, 32 Vassar St, Cambridge, MA 02139; Arizona State University, School of Life Sciences, 427 E Tyler Mall, Tempe, AZ 85281

**Author notes:** Email addresses:* (Jiajia Zhao), (Nancy Lynch), (Stephen C. Pratt).

**Keywords:** Collective Behavior, Decision Making, Ant Colony

## Abstract

The decentralized cognition of animal groups is both a challenging biological problem and a potential basis for bio-inspired design. The understanding of these systems and their application can benefit from modeling and analysis of the underlying algorithms. In this study, we define a modeling framework that can be used to formally represent all components of such algorithms. As an example application of the framework, we adapt to it the much-studied house-hunting algorithm used by emigrating colonies of *Temnothorax* ants to reach consensus on a new nest. We provide a Python simulator that encodes accurate individual behavior rules and produces simulated behaviors consistent with empirical observations, on both the individual and group levels. Critically, through multiple simulated experiments, our results highlight the value of individual sensitivity to site population in ensuring consensus. With the help of this social information, our model successfully reproduces experimental results showing the high cognitive capacity of colonies and their rational time investment during decision-making, and also predicts the pros and cons of social information with regard to the colonies’ ability to avoid and repair splits. Additionally, we use the model to make new predictions about several unstudied aspects of emigration behavior. Our results indicate a more complex relationship between individual behavior and the speed/accuracy trade-off than previously appreciated. The model proved relatively weak at resolving colony divisions among multiple sites, suggesting either limits to the ants’ ability to reach consensus, or an aspect of their behavior not captured in our model. It is our hope that these insights and predictions can inspire further research from both the biology and computer science community.

---

Animal groups are capable of remarkable displays of highly coordinated behavior. Fish schools collectively choose foraging sites (Ward et al., 2012), locusts self-organize into orderly swarms (Yates et al., 2009), oceanic fish assemble in vast migratory shoals (Makris et al., 2009), and social insects perform a host of collective actions including group foraging, construction of complex nests, and adaptive allocation of tasks across the labor force (Charbonneau and Dornhaus, 2015; Detrain and Deneubourg, 2008; Detrain et al., 2019; Oldroyd and Pratt, 2015; Perna and Theraulaz, 2017; Seeley, 1995). How these group actions result from individual behavior remains a major research challenge. Although well-informed leaders may play a role, group organization is typically very decentralized (Detrain and Deneubourg, 2008; Feinerman and Korman, 2017; Seeley, 1995). Coordination emerges from interactions among large numbers of animals acting on limited local information with appropriate decision rules. Connecting individual behavior to group outcomes is too much for unaided intuition, hence mathematical models and agent-based simulations have become useful tools for understanding. In this paper, we present a model for a notable example of decentralized decision-making: nest site selection by ants of the genus *Temnothorax*.

Models, in combination with experimental studies, have already revealed much about these ants, making them a leading model system for collective decision-making (Pratt, 2019). Colonies live in pre-formed cavities such as rock crevices or hollow nuts; if their home is damaged, they are adept at finding candidate new homes, evaluating each site’s quality, and moving the entire colony to the best one. Their decision emerges from the separate efforts of many scouts, each independently recruiting nestmates to the site it has found. Because recruitment is quality-dependent, better sites accumulate ants more rapidly (Mallon et al., 2001). These differences are amplified by a quorum rule under which scouts accelerate recruitment to a site once its population crosses a threshold; the winner of the race to attain a quorum becomes the colony’s choice (Pratt et al., 2002). An agent-based model has shown that this algorithm helps the colony reach consensus on the best site (Pratt et al., 2005). Other models have shown how a colony can make a good choice even when no individual directly compares sites (Masuda et al., 2015a; Robinson et al., 2011), and how individual behavioral strategies optimize speed/accuracy tradeoffs at the colony level (Franks et al., 2013a; Marshall et al., 2006, 2009; Planqué et al., 2007; Pratt and Sumpter, 2006; Sumpter and Pratt, 2009).

Although successful, models of this process have been limited to the simple challenge of choosing between two distinct and equidistant nests in a controlled laboratory environment. Real colonies face more complex scenarios, such as selecting among several sites of varying quality, avoiding splits when candidate nest sites are identical, and resolving colony splits when they occur (Doering and Pratt, 2019; Sasaki and Pratt, 2012). It also remains unclear how the colony maintains high performance with noisy and heterogeneous individuals, and how individuals modify their behavior to account for changes in context or colony state. In addition, a large body of experimental work has uncovered new colony behavior that has yet to be explored in terms of how well our current understanding of the ants’ collective algorithm can explain them. These include the more complex scenarios mentioned above, as well as effects on decision-making of colony size and emigration distance, colony reconnaissance of potential new homes, and the emergence of group-level rationality despite individual-level irrationality (Franks et al., 2008; Sasaki and Pratt, 2011; Stroeymeyt et al., 2010).

To better capture the complexities of nest-site selection, we develop a new, flexible, general model for the analysis and exploration of the underlying behavioral algorithms. We demonstrate the value of the model by reproducing the results of earlier models showing how the ants’ algorithm can account for decision-making and speed/accuracy tradeoffs in simple two-choice experiments (Pratt and Sumpter, 2006; Pratt et al., 2005). We then extend the model to account for more recent empirical observations, including robust decision-making among larger option arrays and rational colony decisions about decision speed (Sasaki and Pratt, 2012; Sasaki et al., 2019). We also explore a proposed novel role for social information, in which ants directly incorporate nestmate presence into their assessment of nest site quality. We use the model to test the effect of such information on a colony’s ability to decide between two identical nests, a context that poses a particular challenge to consensus formation. Finally, we make predictions about the relationship between quorum size and the speed/accuracy tradeoff, and about the ability of a colony to re-unify after dispersal of its members among multiple competing sites.

Our model touches on several aspects of the emergence of collective intelligence in the house hunting process, but many more are yet to be explored. Therefore, an additional goal is to provide a versatile, easy-to-use and maintainable modeling tool that can be used to quantitatively test hypotheses beyond those included in this paper. From a theoretical perspective, our model can serve as a foundation for simpler models that allow rigorous proofs on convergence speed and accuracy.

The rest of the paper is organized as follows: Section 1 defines a general framework that we believe will be useful not only for this algorithm but for other agent-based distributed algorithms as well. Section 2 applies this framework, with designs and interpretations specific to the house hunting context. Section 3 describes our Python implementation of the house hunting simulator, instructions on running simulations, and their scoring goals and metrics. Section 4 validates our model with experimental statistics on individual behaviors and the collective properties of the colony. These validations give us confidence in the accuracy of our model as we proceed into further confirmations of newly observed colony behaviors with limited experimental results, as listed in Section 5. Section 6 showcases simulation results that establish the power of the new social information rule in the colonies’ ability to reach consensus. Next, Section 7 includes new predictions that await experimental test. Finally, Section 8 summarizes our results and proposes possible future research directions.

## 1. Modeling Framework

In this section, we introduce a general modeling “language” that has the potential to be useful for a wide range of applications. In Section 2 we instantiate this language in the context of the house hunting process in ant colonies.

### 1.1. Agent-based Model

Formally, the components below define the entities in the system and their static capabilities. More explanatory text follows after the list.

- **agent-ids**, a set of ids for agents. Each *agent-id* uniquely identifies an agent. We also define **agent-ids**′ to be **agent-ids** ∪{⊥} where ⊥ is a placeholder for “no agent”. In general, we add ′ to a set name to denote the original set with the addition of a default element {⊥}.
- **external-states**, a set of external states an agent might be in. Each element in the set is an *external-state*. In addition, **all-externals** is the set of all mappings from **agent-ids** to **external-states**. Each element of the set is an *all-external*.
- **internal-states**, a set of internal states an agent might be in. Each element in the set is an *internal-state*.
- **env-states**, a set of states that the agents’ environment might take on. Each element in the set is a *env-state*.
- **action-types**, a set of the types of actions agents might perform. Each element in the set is an *action-type*.
- **env-choices**, a set of values an agent can access in the environment. Each element in the set is an *env-choice*.
- **actions**, a set of quadruples of the form (*action-type, agent-id, agent-id′, env-choice*) ∈ **action-types** × **agent-ids** × **agent-ids**′ × **env-choices**. Each element in the set is an *action*.
- **select-action**(agent-id, *state, env-state, all-external*): A *state* is a pair of (*external-state, internal-state*) ∈ **external-states** × **internalstates**. Each (*agent-id, state, env-state, all-external*) quadruple is mapped to a probability distribution over the sample space of **actions**, for which the second component is equal to the input argument *agent-id* and the third component is not equal to it. The function then outputs this probability distribution.
- **transition**(*agent-id, state, all-external, action*): A *state* is a pair of (*external-state, internal-state*) ∈ **external-states** × **internal-states**. Each (*agent-id, state, all-external, action*) quadruple determines a *state* as the resulting state of the agent identified by the input argument *agent-id*. The function outputs the resulting *state*.

Each agent has a unique *agent-id* ∈ **agent-ids**, and is modeled by a state machine. Agents can transition from one *state* to another. A *state* is a pair: an *external-state* ∈ **external-states** that is visible to other agents, and an *internal-state* ∈ **internal-states** that is invisible to other agents.

We define **all-externals** to be the set of all mappings from **agent-ids** to **external-states**. Each element of the set is an *all-external* and represents a particular mapping from **agent-ids** to **external-states** where each *agent-id* is mapped to an *external-state*.

The set **env-states** represents the set of states that the agents’ environment might take on. In this paper, we will assume that the environment is fixed. That is, the *env-state* does not change during the execution of the system. The reason we use a set here is to enable us to model the same set of agents operating in different environments.

Agents can also access values in the environment, and each value is called an *env-choice*. The set **env-choices** is the set of all possible values for *env-choice*.

An agent can transition from one *state* to another by taking an *action* ∈ **actions**. Each *action* consists of an *action-type* ∈ **action-types**, the id of the initiating agent *agent-id* ∈ **agent-ids**, the id of the (optional) received agent *agent-id′* ∈ **agent-ids**′, and *env-choice* ∈ **env-choices**.

The function **select-action**(*agent-id, state, env-state, all-external*) is intended to select an *action* for the agent with the given *agent-id*, who is the initiating agent in the *action*. The function outputs a probability distribution over the sample space **actions**. However the sample space limits its elements to have the second component equal to the input argument *agent-id*, and the third component not equal to it. Thus, any sampled *action* will have *agent-id* being the initiating agent’s id, and the (optional) receiving agent necessarily has a different id.

The function *transition*(*agent-id, state, all-external, action*) represents a transition to be performed by the agent identified by the input argument *agent-id*. Given the input arguments, the function deterministically outputs the resulting *state* of the transition.

### 1.2. Timing and Execution Model

In this section, we introduce the dynamic aspects of our model, including the discrete and synchronous timing model, and how different components in the system interact with each other at different points during the execution of the algorithm.

Our system configuration contains 1) an environment state, called *env-state*, and 2) each agent’s *state*, which is a pair (*external-state, internal-state*), independent of *env-state*. Agents receive inputs from and react to the environment during the execution of the system. In this paper, we will assume that the environment is fixed. That is, the *env-state* does not change during the execution of the system.

Incorporating some theoretical ideas from (Ghaffari et al., 2015; Radeva, 2017), we divide the total time into *rounds*. Each round is a discrete time-step, and times are the points between rounds. At any time *t*, there is a corresponding system configuration *t*. The initial time is time 0, and the first round is round 1, taking the system from configuration 0 at time 0 to configuration 1 at time 1. In general, round *t* starts with system configuration (*t* − 1). During round *t*, agents can perform various **transition**’s, which take the system from configuration (*t* − 1) at time (*t* − 1) to configuration *t* at time *t*.

We now describe the execution of an arbitrary round *t*. At any point in the execution of round *t*, each agent *x* is mapped to a *state, state_x*, which is visible to agent *x* itself. However, to other agents, only agent *x*’s *external_state, external_x* is visible. We denote *all-external* ∈ **all-externals** to be the mapping from every *agent-id* ∈ **agent-ids** to the corresponding *external-state* ∈ **external-states** in round *t*. These mappings can be updated during the execution.

Accounting for the randomness of the order of execution for all the agents, a randomly chosen permutation of **agent-ids** is generated at the beginning of round *t*, serving as the order of execution for the agents in the round. We also instantiate a set 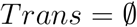 at the beginning of the round. An agent is prevented from changing its *state* further in the round once it adds its *agent-id* to *Trans*, which can happen during its turn (even if there is no resulting state change) or when it performs a **transition** during another agent’s turn. As a result, each agent can change its state at most once in the round. After all agents are in the set *Trans*, round *t* is over, and all agents enter round *t* + 1 synchronously.

The rest of this section describes all possible operations during one agent *x*’s turn in round *t*. When an agent with *agent-id x* (a.k.a. agent *x*) gets its turn to execute, it first checks whether *x* ∈*Trans*. If so, agent *x* does nothing and ends its turn here.

Otherwise, agent *x* has not yet transitioned in round *t*. Let *state_x* denote the *state* of agent *x*. Agent *x* calls the function **select-action**(*x*, *state_x, env-state, all-external*). The function outputs a probability distribution over the sample space of a subspace of **actions**, for which the second component is *x*, and the third component is not *x*. Agent *x* randomly selects an *action*, *act* = (*a, x, x′, e*), according to this probability distribution.

Agent *x* then calls **transition**(*x*, *state_x, all-external, act*), to determine the resulting *state, new_state_x*, for agent *x*. As the initiating agent, *x* also gets added to *Trans*. Next, in the case where *x′* ≠ ⊥, agent *x′* also calls **transition**(*x′, state_x′, all-external, act*) where *state_x′* is the current *state* of agent *x′*, maps itself to the function output, and updates its entry in *all-external*. Note that *x′* is added to *Trans* if the function output is different than *state_x′* in any way. This is the end of agent *x*’s **transition** call. Agent *x* then maps itself to the resulting state *new_state_x*, and updates its entry in *all-external*. Agent *x* finally ends its turn here.

### 1.3. Discussion

Although our model keeps track of the *external-state* of all the agents in *all-external*, when performing a transition, an agent can only access *local* information in it. Locality here is flexible to the context, i.e. local to the location of the agent initiating an action.

Agent-based models are especially powerful for simulating and analyzing collective behaviors given their natural compatibility with object-oriented programming methodologies and their flexibility for allowing individual differences in realized state transition probabilities among the agents (De Vries and Biesmeijer, 1998; Sumpter et al., 2001; Masuda et al., 2015b; Pratt et al., 2005).

## 2. House Hunting Model

This section uses the framework defined in Section 1 to describe the house-hunting process.

### 2.1. Informal Description

An ant colony is composed of **adult workers** and brood items (immature ants), each group making up 40% to 60% of colony members. Adults are roughly equally divided between active workers, who organize and execute emigrations, and passive workers, who, like brood items, are typically transported to the new nest by active workers and not themselves recruit nestmates (Pratt et al., 2005; Dornhaus et al., 2008; Valentini et al., 2020a).

There are four distinct phases for an active worker in the house-hunting process. In the first, the **Exploration** phase, the ant randomly starts to explore her surroundings for a suitable new nest. If she finds a candidate site, she enters the **Assessment** phase, where she individually assesses the site’s quality according to various metrics (Healey and Pratt, 2008; Franks et al., 2003b; Pratt, 2005a). If she judges the site to be satisfactory, the ant accepts it and enters the **Canvassing** phase, in which she returns to the old nest to recruit other ants to the site by leading **forward tandem runs** (FTR). In a FTR, the recruiter slowly leads a single follower (another active worker) from the old nest to the new (Moglich, 1978; Richardson et al., 2007; Valentini et al., 2020b). Upon arriving at the nest, the follower ant goes directly into the Assessment phase and evaluates the nest’s quality independently of the leader ant. If she finds the nest satisfactory, she will transition to the Canvassing phase and start leading FTRs to the nest. A canvasser continues leading FTRs until she perceives that the new nest’s population has exceeded a threshold, or quorum (Pratt, 2005c). At this point, she enters the **Transport** phase, in which she fully commits to the new nest as the colony’s home. She ceases FTRs and instead switches to picking up and carrying nestmates from the old to the new nest. These transports are faster than FTRs, and they are largely directed at the passive workers and brood items, hence they serve to quickly move the entire colony to the new nest (Pratt et al., 2002, 2005). Previous models and experiments indicate that the quorum rule helps the colony to reach consensus rather than splitting among multiple sites (Pratt et al., 2002; Franks et al., 2006, 2013b). Splitting becomes a danger if ants at different sites, each ignorant of their nestmate’s discoveries, launch FTRs to their respective sites. The quorum rule makes it likely that whichever site first hits the threshold will quickly end up with all or most of the colony, due to the speediness of transport.

Although experimental evidence is equivocal, we assume that the quorum size is correlated with the number of adult workers in the colony (Dornhaus and Franks, 2006; Franks et al., 2006). We also assume that passive workers can contribute to quorum attainment. Once the quorum is met, the switch to Transport phase is irreversible: an ant continues transporting nestmates to her new home nest even if the nest population later drops below the quorum size (Pratt, 2005c). However, transporters do sometimes interrupt transport to search for and assess alternative nest sites. If the search yields a new site that is better than the ant’s current nest, then she exits the Transport phase and enters the Assessment phase with the new site as her candidate nest.

An ant in the Canvassing or Transport phase does not recruit indefinitely. Once the site from which she is recruiting is empty, she returns to her home nest and transitions back to the **Exploration** phase. However, this happens only upon meeting a “termination” condition consisting of ten occurrences of either of the following events: 1) the worker tries to lead a FTR where the solicited follower is also trying to lead her own FTR, and 2) the worker tries to carry another worker who is also in the Transport phase. This condition is based on frequent observation of these events at recently emptied nests. We hypothesize that an ant’s requirement of several such events is a means of ensuring thorough exploration of the old nest so that no nestmates are left behind. We do not have a precise measure of how many such events are required, but chose the number 10 as an upper-bound estimate.

The emigration is completed when all ants in the colony are relocated to the new nest, except possibly for a few active scouts (Pratt et al., 2002).

### 2.2. Formal Model

#### 2.2.1. Model components

In this section, we show how each component in our modeling framework (Section 1.1) is defined in the house hunting algorithm context.

Fig. 1 shows our native data structures as used by various components in the system: *Nest* objects, an array which constitutes an *env-state*; *Ant* objects, each corresponding to an agent; *State* = (*External-State, Internal-State*)objects, each corresponding to a *state* = (*external-state, internal-state*), and *Action* objects, each corresponding to an *action*. Each of the data structures contains a set of variables, as seen in Fig. 1. Note that we consider all variables belonging to either the class *ExternalState* or the class *InternalState* to belong to the class *State* as well. Throughout the rest of the paper, we use the notation *object.variable* to denote the value of a *variable* belonging to a class *object*. Using these data structures as building blocks, we now show all possible values for the components in the framework presented in Section 1.1. Note that for consistency with our implementation in Section 3, we use −1 or an empty string “” to represent any invalid default integer or string values represented by ⊥ in Section 1.

**Figure 1:**
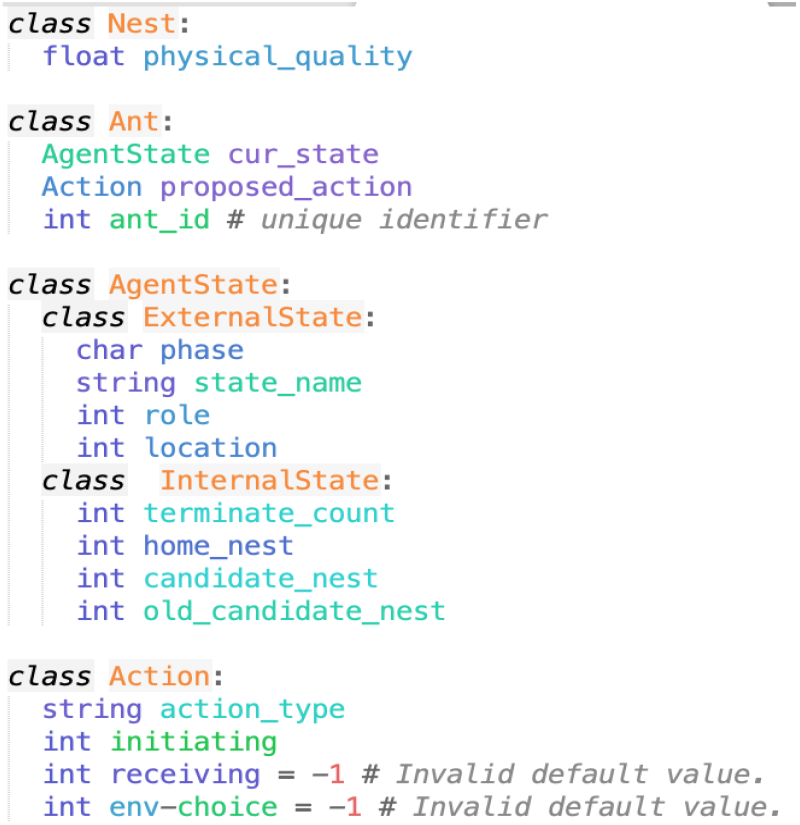
Native data structures that define different entities in the distributed system.

- **agent-ids**, the set containing all integers in the range [0, num_ants), where num_ants is the total number of ants in the colony. In addition, **agent-ids**′ = **agent-ids** ∪ −1. Each *Ant* is initialized with its corresponding *ant_id*, which corresponds to a *agent-id*.
- **external-states**, the set containing all possible values for an *External-State* class object, each corresponding to an *external-state*. We designed these variables to be in the *external-state* because these contain information that influences other ants’ activities. Therefore, it is biologically plausible that individuals have access to this information about one another. In any *ExternalState* class object, *phase* has one of four possibilities - Exploration (searching for new nests), Assessment (assessing new nests), Canvassing (leading other active workers on FTRs to her accepted candidate nest), and Transport (committing to the new nest and rapidly carrying other ants to it). Note we abbreviate the four phases to names “E”, “A”, “C” and “T”, respectively. The initialization of an *Ant*’s *phase* and *state_name* can be found in Section 2.2.3. For each *phase*, the variable *state_name* take values from a different set, as follows:

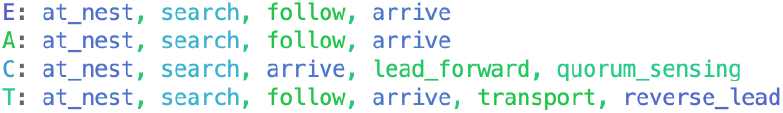 The variable *role* can be one of (0,1,2) representing (active ant, passive ant, brood), and each *Ant* is initialized with the appropriate value. The variable *location* can be any integer in the range [0, num_nests) where num_nests is the total number of nests in the environment, with 0 representing the original home nest. In addition, recall that **all-externals** is the set of all possible mappings from **agent-ids** to **external-states**. Each element of the set is an *all-external*.
- **internal-states**, the set containing all possible values for an *Internal-State* class object, each corresponding to an *internal-state*. The set of fields we designed for the *InternalState* class represent information that should only be accessed and modified by an ant’s internal memory. Each of *home_nest* (initial value = 0), *candidate_nest* (initial value = −1), and *old_candidate_nest* (initial value = −1) can take any integer in the range [0, num_nests), where num_nests is the total number of nests. Lastly, *terminate_count* (initial value = 0) takes any value in the range [0, 10].
- **env-states**, a set of arrays, each being an array of the *Nest* class objects. Each array corresponds to an *env-state*. For an *env-state*, the *Nest* at index 0 represents the original home nest and has *physical-quality* 0. All other *Nest*’s have *physical-quality* in range [0, 4]. The maximum quality 4 here is arbitrary. Recall that the array does not change throughout the execution of the system, and the array is read from a configuration file introduced in Section 3.1.
- **action-types**, the set of the types of actions includes: “search”, “no_action”, “find”, “follow_find”, “get_lost”, “reject”, “no_reject”, “accept”, “recruit”, “quorum._met”, “quorum.mot_met”, “stop_trans”, “delay”, “terminate”, “lead”, “carry”. *Action-type* is initialized to “no_action”. Each item in the set above is an *action-type*.
- **env-choices**, the set of integers in [0, num_nests) ∪ −1 where num_nests is the number of nests in the environment. Each element in the set is an *env-choice* and is an integer representing an index into *env-state*. An *env-choice* has initial value −1.
- **actions**, the same set as defined in Section 1.1. Note that not all actions require a receiving agent, and not all actions require an *env-choice*. In case that they are not needed, they take the invalid default value −1.
- **select-action**(*agent-id, state, env-state, all-external*): the same function as defined in Section 1.1. Refer to Section 2.2.2 for details.
- **transition**(*agent-id, state, all-external, action*): the same function as defined in Section 1.1. Refer to Section 2.2.3 for details.

#### 2.2.2. The select-action function

The function **select-action**(*x, state_x, env-state, all-external*) outputs a probability distribution over the sample space of **actions**, for which the second component is equal to the input argument *agent-id* and the third component is not equal to it. Let any *action* in the sample space be denoted by (*a, x, x′, ec*), where the second component is fixed. We now list out the probability distribution on other components for each possible value of the *state_name* variable in *state_x*, as it is the only variable in *state_x* that affects the output probability distribution. The boldface words are parameters that we can tune and whose values are read from a configuration file, introduced in Section 3.1.

- For *search*, the probabilities of choosing *a* to be “find” and “no_action” are **search_flnd** and 1-**search_find** respectively, and all other *action-type*‘s have 0 probability. Both variables *x′* and *ec* take the invalid default value −1 with probability 1.
- For *follow*, the probabilities of choosing *a* to be “follow_find” and “get_lost” are **follow_find** and 1-**follow_flnd** respectively, and all other *action-type*’s have 0 probability. Both variables *x′* and *ec* take the invalid default value −1 with probability 1.
- For *reverse_lead*, the probabilities of choosing *a* to be”delay” and “no_action” are **transport** and 1-**transport** respectively, and all other *action-type*’s have 0 probability. Both variables *x′* and *ec* take the invalid default value −1 with probability 1.
- For *quorum_sensing*, let the set 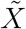 be the set containing id’s of all agents with *external-state* having *role* ∈ {0, 1}and *location* = *state_x.location*. If the set size 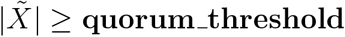, the probabilities of choosing *a* to be “quorum_met” and “quorum._not._met” are 1 and 0 respectively, and are 0 and 1 otherwise, and all other *action-type*’s have 0 probability. Both variables *x′* and *ec* take the invalid default value −1 with probability 1.
- For *leadforward*, let 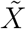 be the set containing id’s of the agents that are not *x*, and whose *external-state* has *role* = 0 and *location* = *state_x.location*. The function selects an action 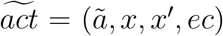 according to the following probability distribution. In case *terminate_count* < 10, 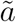 is chosen among “lead” and “get_lost” with probabilities **lead_forward** and 1-**lead_forward** respectively, and all other *action-type*’s have probability 0. In case *terminate_count* ≥ 10, 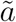 is “terminate” with probability 1. The variable *ec* is equal to {*state_x.candidate_nest*} with probability 1. The distribution of *x′* depends on 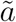, as follows:
  – For “lead”, if 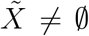, the variable *x′* is uniformly selected from 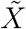, and all other values in *agent-id′* have 0 probability; otherwise, *x′* = −1 with probability 1.
  – For “get_lost”, *x′* = −1 with probability 1.
  – For “terminate”, *x′* = −1 with probability 1.
- For *transport*, let 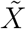 be the set containing id’s of all agents that are not *x*, and whose *external-state* has *location* = *state_x.location*. In addition, let 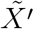 be the subset of 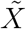 containing agents that have *role* ∈ {1, 2}. The function first selects an action 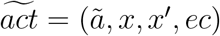 according to the following probability distribution. In case *terminate_count* < 10, 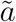 is chosen among “carry” and “stop_trans” with probabilities **transport** and 1-**transport** respectively, and all other *action-type*s have probability 0. In case *terminate_count* ≥ 10, 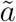 is “terminate” with probability 1. The variable *ec* is equal to { state _x.home_nest} with probability 1. The distribution of *x′* depends on 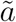, as follows:
  – For “carry”, if 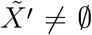, *x′* is uniformly sampled from 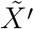, and all other values in *agent-id′* have 0 probability. Otherwise if 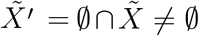, *x′* is uniformly sampled from 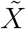, and all other values in *agent-id′* have 0 probability. Otherwise, *x′* = −1 with probability 1.
  – For “stop_trans”, *x′* = −1 with probability 1.
  – For “terminate”, *x′* = −1 with probability 1.
- For *at_nest*, the probability of choosing *a* to be “search” is 1 − *p*(*x*), where *x* is the quality of the nest option under assessment (Figure. 1) and *p*(*x*) defined in Equation 2. There are always two possible actions for a *state* with *state_name* = *at_nest*, and the one that is not “search” naturally has probability *p*(*x*). All other *action-type*’s have 0 probability. Both variables *x′* and *ec* take the invalid default value −1 with probability 1. To determine *p*(*x*), an ant is required to assess the quality of a nest in the environment. The assessment of the quality of a nest includes both its physical qualities (Burns et al., 2016; Sasaki et al., 2015) and the nest population (Pratt, 2005d; Dornhaus and Franks, 2006). Therefore, we use a simple linear combination of these two values to denote the final nest quality with a new parameter called **pop_coeff** as the coefficient of the population effect. In other words, the final nest quality of a nest with *physical_quality q* and population *pop* (obtained from *all-external*) is

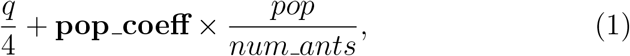

where 4 is the maximum value of nest qualities, and *num_ants* is the total colony size. We further define the following sigmoidal function (Fig. 2)

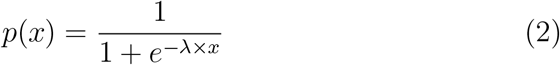

where λ is a parameter that controls how “steep” the sigmoidal function is, and *x* is the above defined nest quality. Higher λ values correspond to lower individual noise level, and bring *p*(*x*) closer to a step function.
- For *arrive*, the probabilities of choosing *a* to be “reject” and “no_reject” are 1 − *p*(*x*) and *p*(*x*) respectively, where *x* is the difference in quality of the candidate nest compared to the home nest (Equation 3) and *p*(*x*) defined in Equation 2. All other *action-type*’s have 0 probability. The variable *x′* take the invalid default value −1 with probability 1. The variable *ec* take the invalid default value −1 with probability 1. To determine *p*(*x*), an ant *x* is required to compare the quality of its *candidate_nest* (with physical_quality *q*_1_ and population *pop*_1_) and its *home_nest* (with physical_quality *q*_0_ and population *pop*_0_). We still use the sigmoidal function in Equation 2, with the change that the input *x* to the function now is

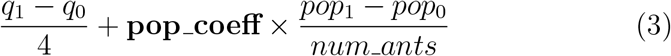

where 4 is the maximum value of nest qualities, and *num_ants* is the total colony size.

**Figure 2:**
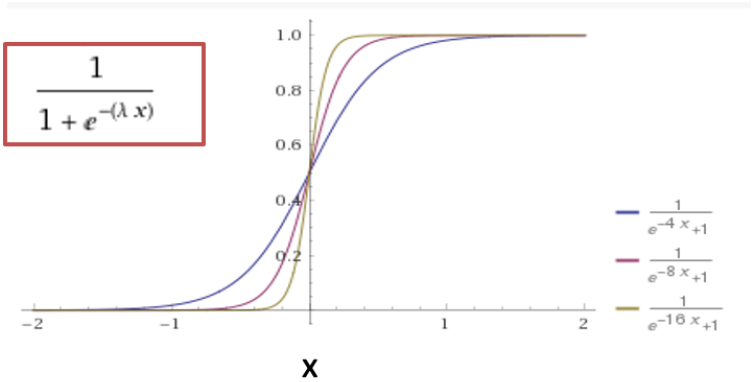
Sigmoidal function with λ = 4, 8, 16.

#### 2.2.3. The transition function

##### Passive Workers and Brood Items

Active worker scouts are defined as those who engage in the emigration process by independently discovering the new nests (entering without carrying or being carried) or by carrying brood items or other adult ants to the new nest or both. Passive workers remain in the old nest until they are carried to the new nest. Brood items are similar to passive workers but do not contribute to quorum attainment (Pratt et al., 2002; Dornhaus et al., 2008).

We use *sn_p_* to denote a *state* with a certain *state_name* = *sn* and *phase* = *p*. Passive workers and brood items together form the passive majority population in the colony. Their behavior pattern is thus very simple — they only have one *state_name_phase_, at_nest_E_*, available to them. They can only allow one *action* with *action-type* “carry” and themselves as the receiving agent. The action results in the *location* variable in their *state* set to the last component of the action, *env-choice*, and no other variables in their *state*’s can change throughout the execution. Therefore, the rest of the section focus on the *state* transitions of **active** workers only, including any initiating and receiving ants involved.

##### Initiation and Termination of Emigration

All ants start in *at_nest_E_*. Their *role* variable values are assigned the corresponding numbers, and *home_nest, location* are both initiated with 0, the original home nest. The variables *candidate_nest* and *old_candidate_nest* are set to −1 as the default invalid value. And *terminate_count* starts with 0.

We do not designate a separate “termination state” that disables an ant from exploring further, but at the termination of the emigration process, we expect most active workers to be in *at_nest_E_*. This is enforced softly through the population effect introduced in Section 2.2.2 - if an agent in *at_nest_E_* is in a nest with both a high physical quality and a high nest population it is highly likely that she is happy staying put in this nest and stabilizes in the state *at_nest_E_*. As a result, the more agents stabilizes in the same nest, the more likely that they will stay stable and that new agents will stabilize as well. In the house hunting algorithm, the conditions that trigger this “termination” behavior contains two cases, as mentioned in Section 2.1. The details of this special “termination” case handling is discussed in the next paragraph.

##### Special and General Cases

In the house-hunting algorithm, there are some special cases that the **transition** function handles before outputting the resulting *state*. To facilitate, we define a set **allowed-in**(external-state) to be a mapping from **external-states** to subsets of **action-types**. Consider an *external-state s*, and the allowed subset is then **allowed-in**(*s*), representing the set of actions the agent in the *external-state s* is allowed to receive. The four variables *s* contains (as shown in the *ExternalState* class) each affects **allowed-in**(*s*) in the following way. *location* has no influence. If *role* is 1 (passive) or 2 (brood), **allowed-in**(*s*) = “carry”. Otherwise, *role* = 0. Let *state_name_phase_* denote the *state_name* and *phase* variables in *s*. For *at_nest_E_*, *at_nest_A_*, and *at_nest_T_*, **allowed-in**(*s*) = “lead”, “carry”. For *search_E_*, *search_A_, search_C_*, *search_T_*, and *at_nest_C_*, **allowed-in**(*s*) = “carry”. For all other cases, 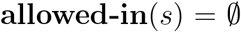.

We now list out how the function *transition(agent-id, state, all-external*, *action*) handles each of the special cases, and also the general case. Let the input argument *action* be expanded to the quadruple (*act = a, x, x′, ec*). Also recall that the set *Trans* is a set containing the id’s of all the agents that have completed a *state* change in the round (Section 1.2).

- The first special case is if the input argument *agent-id* = *x′*. This case only happens when agent *x′* ≠ −1 invokes (in agent *x*’s turn) a **transition**(*x′*, *state, all-external, act*), where *state* = (*external′, internal′*) is the current *state* of agent *x′*. If *x′* ∈ *Trans* or if *a* ∉ **allowed-in**(*external′*), the function simply ends by returning the input argument *state*. Otherwise, the function adds *x′* to *Trans*. It then finds the black text box corresponding to *state.phase* and *state.state_name* in Fig. 3a, and the black text box that *a* leads to contains the *phase* and *state_name* of the resulting *state*. The rest of the variables in *state* are modified as well, and the details are listed for each possible value of the (*phase, state_name*) pair at the end of the section. The function then outputs the resulting *state*.
- The second special case is if *act* satisfies the termination condition mentioned earlier in this Section. Specifically, the cases are when *agent-id* = *x*, and *act* is either 1) (*lead_forward, x, x′, state_x.candidate_nest*) and *x′* = ≠ −1 has an *external-state* with *state_name* = *lead_forward*,or 2) (*transport, x, x′, state_x.home_nest*) and *x′* ≠ −1 has an *external-state* with *state_name* = *transport*. We call these the “termination conditions”. When *act* satisfied either clauses, after adding *x* to *Trans*, the function ends its execution by outputting a resulting *state* that only differs from the input argument *state* by adding 1 to the *terminate_count* variable.
- The third special case is if *agent-id* = *x* and *act* does not satisfy the termination conditions, but *x′* ≠ −1 and either of the following is true: 1) *x′* ≠ *Trans*, or 2) *a* ∉ **allowed-in**(*external′*). Note the second case here excludes cases that satisfy our termination conditions stated in the last bullet point. In other words, the second special case has priority over this third special case. In this third special case, the function adds *x* to *Trans*, and ends its execution by outputting the original input argument, *state*.
- Lastly, in the general case where none of the above special cases applies, the function first adds *x* to *Trans*. Then it finds the black text box corresponding to *state.phase* and *state.state_name* in Fig. 3b, and the black text box that *a* leads to contains the *phase* and *state_name* of the resulting *state*. The rest of the variables in *state* are modified as well, and the details are listed for each possible value of the (*phase, state_name*) pair at the end of the section. The function then outputs the resulting *state*.

**Figure 3:**
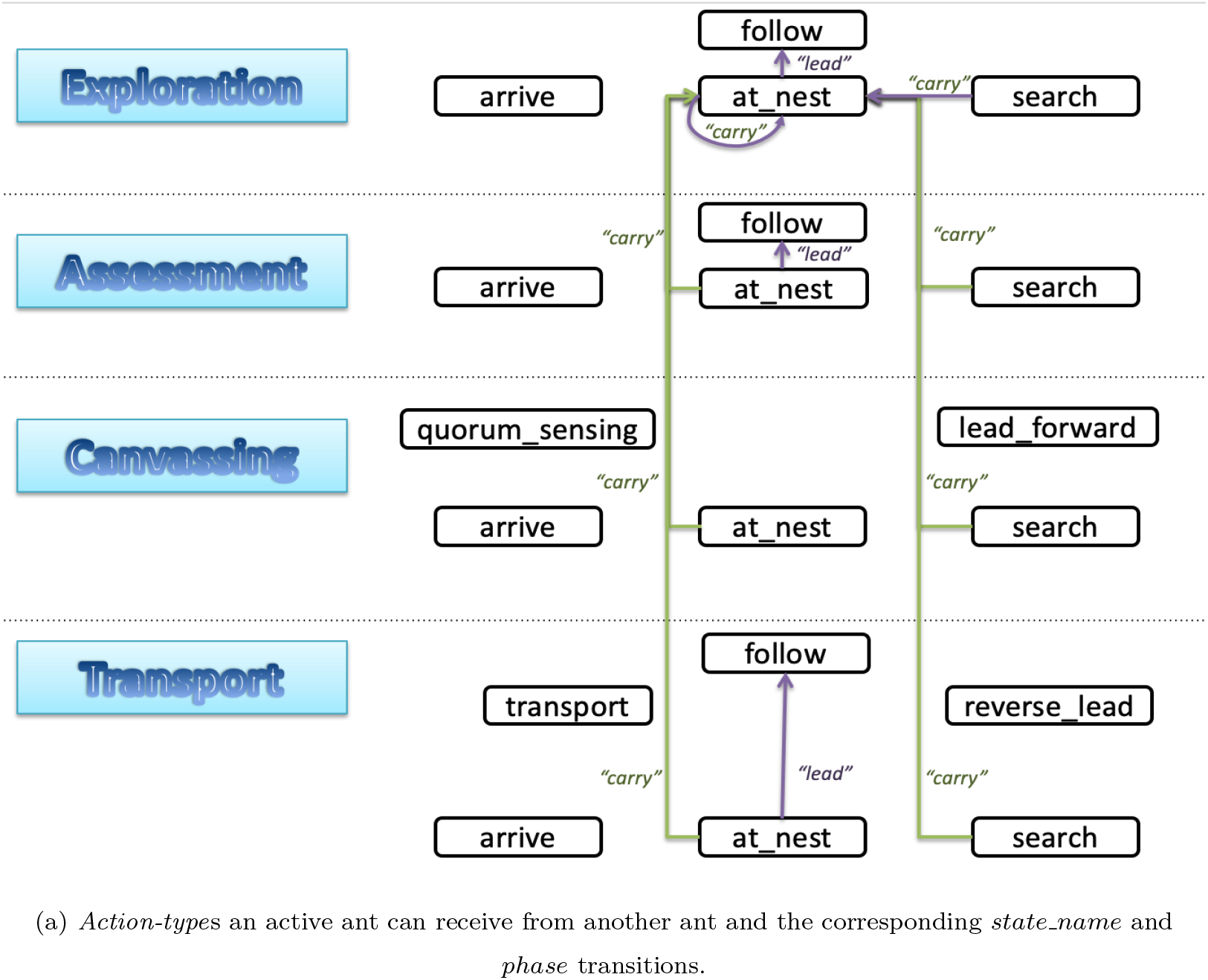
States and actions modeling the behavior of active ants responsible for organizing colony emigrations. As described in Section 2.1, the four distinct phases are in different boxes: Exploration, Assessment, Canvassing, and Transport.

**Figure 3:**
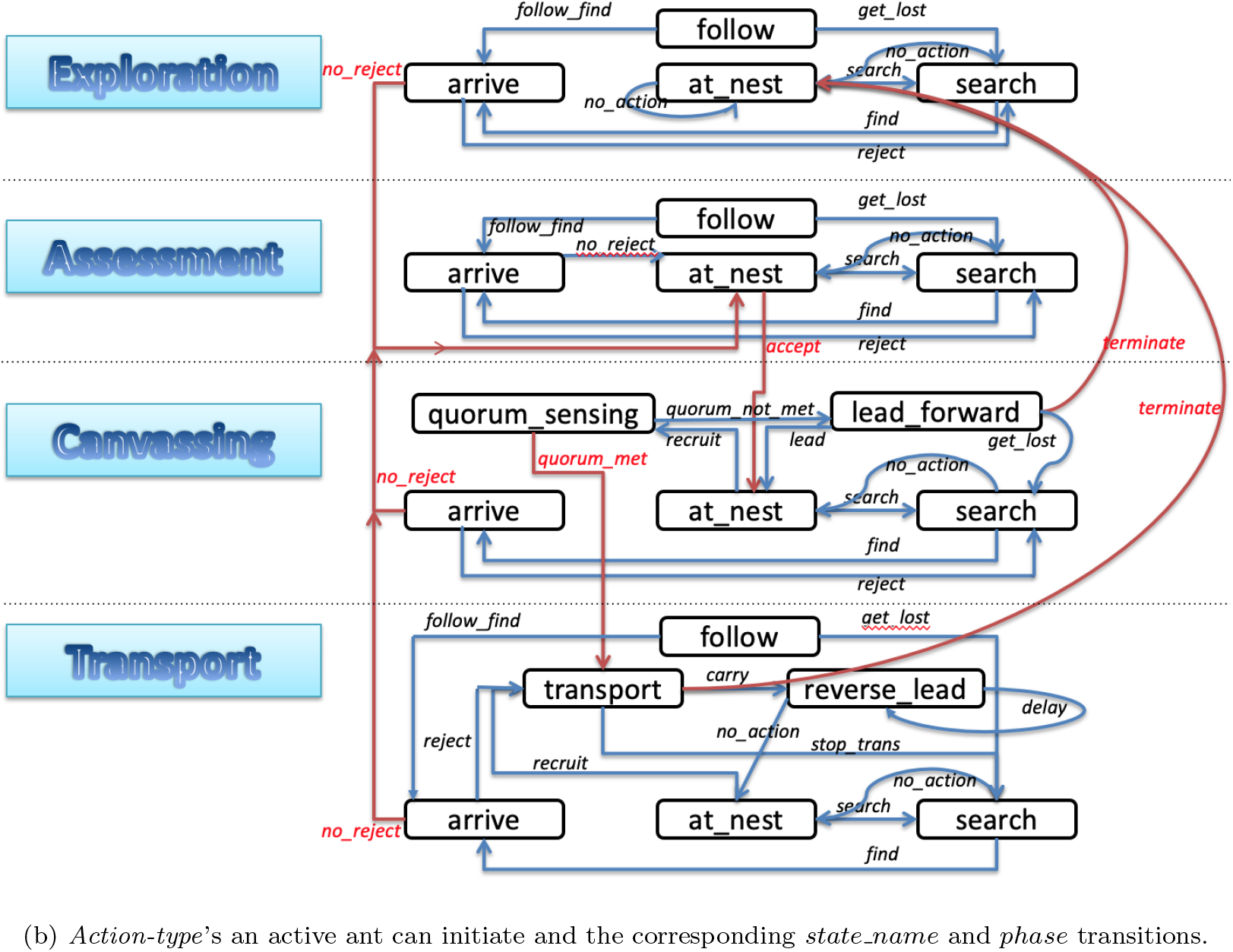
States and actions modeling the behavior of active ants responsible for organizing colony emigrations. As described in Section 2.1, the four distinct phases are in different boxes - Exploration, Assessment, Canvassing, and Transport.

In Fig. 3a and Fig. 3b, *action-type*’s are color-coded as shown in Table 1. We walk through in the supplement materials all possible transitions of an ant and the associated changes in the internal and external states. An example is an ant in *at_nest_E_* has four possible actions. First, she can perform “no_action” and remain in the current nest. Second, she can perform “search” and go into the state *search_E_*. Third, she can receive a “lead” by another ant to follow a FTR to a destination nest, *ec* ∈ **env-choices**, in which case she sets *old_candidate_nest* to the value of *candidate_nest*, and sets *candidate_nest* to *ec*. Then she transitions to the state *follow_E_*. Finally, she can receive a “carry” by another active worker ant to a destination nest *ec* ∈ **env-choices**, in which case her *location* and *candidate_nest* are changed to ec, and she stays in *at_nest_E_*.

**Table 1:** Color coding of arrows representing *action-type*’s in Fig. 3a and Fig. 3b.

| Arrow Color | Init/Recv | Phase Change |
| --- | --- | --- |
| blue | Initiating | No |
| red | Initiating | Yes |
| purple | Receiving | No |
| green | Receiving | Yes |

## 3. Model Simulation and Metrics

In this section we describe the configuration file that contains all the parameters of the model in Section 2. We also quantitatively define the speed and accuracy measures for our simulation runs. A detailed description of the Python simulator implementation is in the supplemental material.

### 3.1. Configuration Parameters

There are three kinds of parameters: environment, algorithm, and settings.

Environment parameters are controlled by the environment and not considered changable or tune-able. These include the number of ants in the colony, and the number and physical qualities of the nests as potential new nest options.

Algorithm parameters are parameters that we can manipulate in order to change the *select-action* function and hence the outcomes of our simulations. These include the λ for the sigmoid function in Equation 2, the pop_coeff value, parameters related to quorum sizes, the probability of finding a new nest in the environment, the probabilities of following and leading a FTR without getting lost, and the probability of continued transports instead of stopping transportation. Related explanations are in Section 2.2 and 2.2.2.

Settings parameters control plotting features and also convergence criteria. These include the option to generate a plot, the total number of runs for every environment/algorithm setting, the maximum number of rounds per simulation run, the percentage of ants needed in a nest to declare **convergence**, and the number of rounds the convergence needs to persist to declare **persisted convergence** which marks the end of the simulation run. An example is in the supplemental materials.

#### Baseline Default Parameter Values

Compared to the agent-based model in (Pratt et al., 2005), our model places less emphasis on assigning specific observed values to a large number of parameters, but rather on a simple and elegant model that is more agile in representing a wide range of possible behaviors. For that reason, some parameters cannot be directly drawn from existing empirical data. We estimate these parameter values in a trial-and-error fashion until simulation results match well with the empirical results in (Pratt et al., 2005). These baseline values are used as a default from Section 4 to 7, unless otherwise specified.

The sources for determining the parameter values are listed in Table 2. In particular, the values of **lambda_sigmoid**(range: 1 to 16) and **pop_coeff** (range: 0 to 1) are picked by trial-and-error to model individual sensitivity to nest qualities, and the significance of colony information versus individual judgements. The quorum size (**quorum_thre** × (num_active+num_passive) +**quorum_offset**) is observed to have a median value between 4 and 18 ants for worker populations from 24 to 150, with the quorum size having a signif-icant positive correlation with the number of adult ants (Pratt et al., 2002; Franks et al., 2015). Therefore, with a colony of 200 members (including 100 adult workers), we use a **quorum_thre** of 15% and set **quorum_offset** to 0, estimating a quorum size of 15. The value of **search_flncl** (range: 0 to 1) is determined experimentally by trial-and-error. This parameter can be influenced by many other factors such as the spatial geometry of the nests and the experience level of the individual. These nuances are not captured in our model in the interest of simplicity. But they can significantly affect the simulation outcomes, and are an important future extension of our work. The parameter **follow_find** denotes the success rate of a tandem run without the follower getting lost and starting a new search. A successful tandem run requires that both ants reach the target nest. Empirical observations suggest large variation in tandem success, with observed success ranging from 30% to over 90% (Pratt, 2008; Glaser and Grüter, 2018). However, even lost followers enjoy a significantly increased chance of finding the target nest on their own (Pratt, 2008). We thus chose a high FTR success rate of 0.9 to capture both these direct and indirect effects of tandem following on nest discovery. The parameter **lead_forward**(range: 0 to 1) is the probability that an ant performs an FTR when in the *lead_forward* state. The alternative option, *get_lost*, is designed to model the slower speed of an FTR, and is determined experimentally in a trial-and-error fashion. The parameter **transport** is the probability that an ant keeps transporting instead of stopping to resume search for additional sites. The stopping probability is observed to be between 0.06 and 0.44, meaning our **transport** should take values between 0.56 and 0.94. We chose 0.7 as our baseline value.

**Table 2:** Default parameter values and the sources that helped determine these values.

| Parameter | Value | Source |
| --- | --- | --- |
| **lambda_sigmoid** | 8 | trial-and-error, Sec. 6.1 |
| **pop_coeff** | 0.35 | trial-and-error, Sec. 6.1 |
| **quorum_thre** | 0.15 | (Pratt et al., 2002; Franks et al., 2015) |
| **quorum_offset** | 0 | (Pratt et al., 2002) |
| **search_find** | 0.005 | trial-and-error |
| **follow_find** | 0.9 | (Glaser and Grüter, 2018; Pratt, 2008) |
| **lead_forward** | 0.6 | trial-and-error |
| **transport** | 0.7 | (Pratt et al., 2005) |

An average colony size of 200 with 50 active workers, 50 passive workers, and 100 brood items is within the range of real colony compositions (Franks et al., 2015). One round approximately translates to 0.5-1 minutes, though this is a very rough estimate. A simulation with 2000 rounds thus translates to 16-32 hours, and one with 4000 rounds translates to 32-64 hours. The values for variables **percent_conv** and **persist_rounds** are determined by trial-and-error and rough estimates from past empirical observations.

### 3.2. Speed and Accuracy Measures

We define the speed and accuracy metrics below for the *whole emigration process* until either convergence or the end of simulation, including cases resulting in splitting.

#### Convergence Score as Speed

The final goal of the house hunting algorithm is to achieve fast convergence in any given environment and stabilize at that convergence. To assess how well this was achieved, we calculate a **convergence score** as the inverse of the round number when a persistent convergence started. If no persistent convergence was reached before the end of the simulation, the convergence score is 0. Each simulation run has a convergence score.

#### Accuracy

Another important metric is **accuracy**, which is defined for a group of simulation runs. This metric tells us how good the colony is at selecting the best choice in the environment. Thus, each of the nest options in the configuration has an empirical probability of the colony converging to it, called the nest’s **convergence probability**. Note that we also get a probability of splitting. To calculate the final accuracy, we also normalize the nests’ physical qualities, such that the best nest has quality 1 after normalization, and the worst nest, which is the home nest, has quality 0. The **accuracy** of the configuration is then

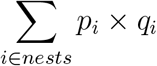

where *p_i_* is nest *i*’s convergence probability, and *q_i_* is its normalized physical quality. If no convergence is reached (splitting), the physical quality corresponding to that probability will be 0, thus not contributing to the summation above.

## 4. Model Validation

We begin by validating our model against the same empirical data that were successfully accounted for by two earlier models (Pratt et al., 2005; Pratt and Sumpter, 2006). First, we examine a simple scenario where colonies have only one candidate nest in the environment. Then we consider a decision between two nests that clearly differ in quality. Finally, we investigate how colonies trade off speed and accuracy depending on the urgency of their move. For all scenarios, we simulate the same data explored by the earlier models, and compare our results, at both individual and colony level, to the empirical observations. All simulations for the rest of the paper default to the configuration file described in Section 3.1, unless specified otherwise.

### 4.1. Single-Nest Emigrations

The first question we ask is: does our model accurately reproduce statistics on individual recruitment acts in single-nest emigrations? Previous empirical work showed the distributions across ants of key behaviors contributing to the collective outcome (Pratt et al., 2005). These include the number of recruitment acts per ant, the number of ants performing each recruitment type, and the number of ants arriving at the new site by different routes. We asked whether our model could replicate the empirical distributions. To answer the question, we simulated the single-nest experiments conducted in (Pratt et al., 2005), on the six colonies with compositions detailed in Table 3. We used default parameter values, except we increased *search_find* to 0.05. This increase accounts for the presence of only one new nest, hence all “find” actions after the first one are re-discoveries of this nest, which we assume has a higher probability than finding a previously unknown site (Pratt et al., 2005). In future work, this variable should be expanded to depend on other factors, such as the number of nests in the environment or the spatial geometry. We ran 500 simulations for each colony.

**Table 3:** Compositions of colonies used in two-nest emigrations for model validation as shown in (Pratt et al., 2005).

| Colony | Active | Passive | Brood | Total |
| --- | --- | --- | --- | --- |
| A4 | 70 | 28 | 228 | 326 |
| A6 | 59 | 74 | 111 | 244 |
| A8 | 62 | 95 | 106 | 263 |
| A14 | 67 | 42 | 192 | 301 |
| A16 | 53 | 88 | 61 | 202 |
| A17 | 73 | 101 | 173 | 347 |

#### Results

We compared the statistics of the model output to the same statistics reported in (Pratt et al., 2005) (Fig. 4). Fig. 4(a) shows the histograms of individual workers grouped by the number of recruitment acts. More than half of the simulated workers never recruited, consistent with the empirical finding of about 60% non-recruiting active workers. The other bins show similar mean and variance to the empirical data. Fig. 4(b) classifies ants by their recruitment behavior, and the breakdowns are again consistent with the experimental observations. Fig. 4(c) categorizes workers by their routes to discovery of the candidate nest, and is again consistent with the findings in (Pratt et al., 2005), at least when the experimental data are pooled over six emigrations by three colonies. However, the distributions across the three different routes vary strongly across emigrations. Indeed, the results in (Pratt et al., 2005) notably differ from those in (Pratt, 2005b). While our model does not account for this variation, we conclude that it does adequately reproduce key distributions in recruitment behavior in single-nest emigrations.

**Figure 4:**
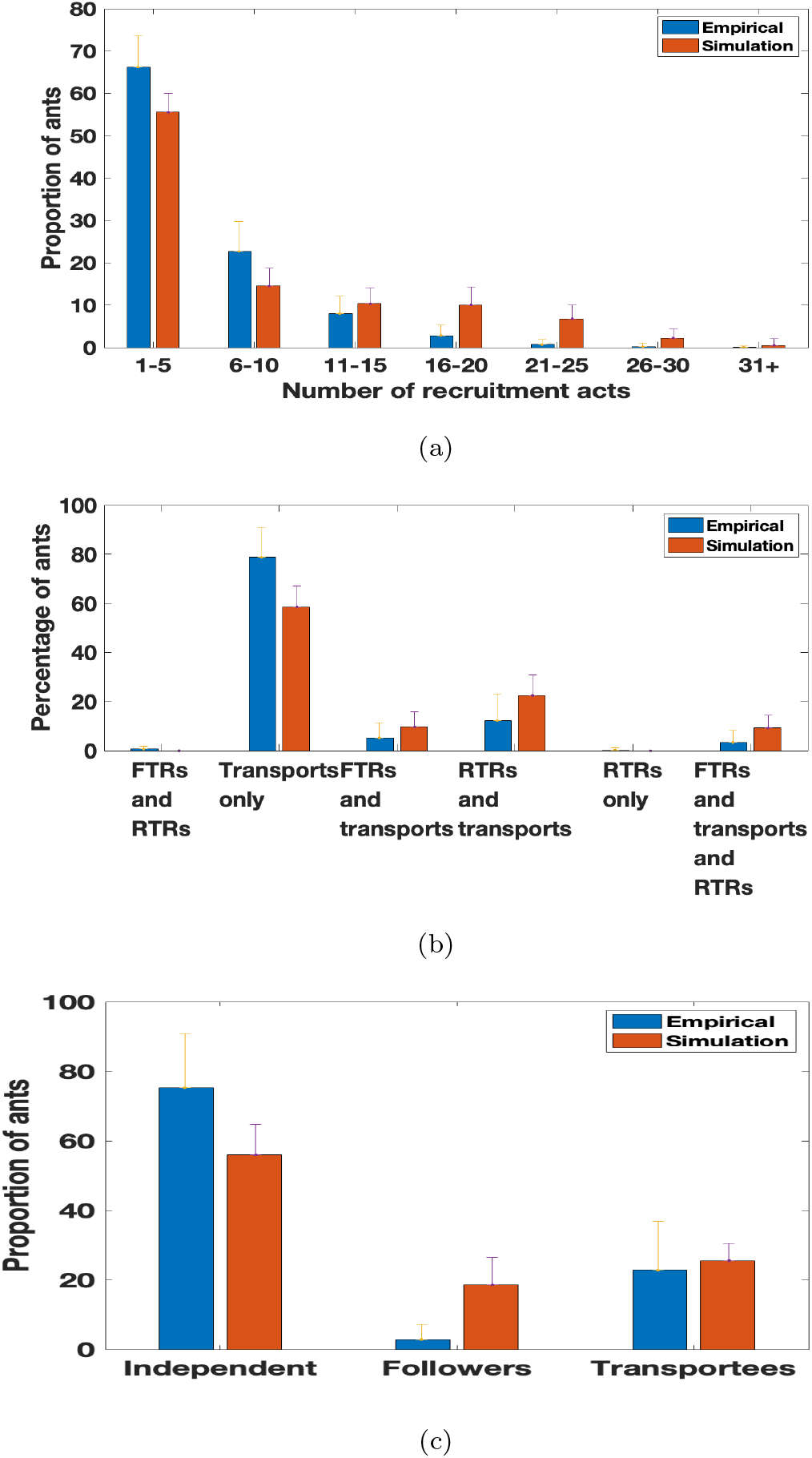
(a): Histogram of workers grouped by the number of recruitment acts performed. (b): Histogram of workers who performed different types of recruitment acts. (c): Histogram of workers grouped by the route by which workers arrived at the new nest. Blue bars are results from (Pratt et al., 2005). Orange bars show our simulation results. Bar values are averaged over 500 simulations. Error bars show standard deviations.

### 4.2. Two Unequal Nest: Splits

The second question we ask is: does our model account for the degree of splitting in two-nest emigrations with unequal qualities? In these circumstances, colonies do not always make a unanimous choice, but may temporarily split between the sites before eventually coalescing on a single one. We focus on splitting because it is a primary hindrance to consensus. We measure splitting as the percentage of brood items in the better candidate nest at the time when the last ant has been moved from the home nest.

We replicated the two-nest emigrations in (Pratt et al., 2005), with six colonies whose member compositions are listed in Table 3. We set **nest_qualities** = [0,1,2], representing a destroyed old nest and two candidate nests of mediocre and good quality, respectively. The rest of the configuration parameters were left at the default values.

We ran 500 simulations for each colony, and for each colony we recorded the average percentage of brood items in the better nest at the time the home nest became empty. To compare the simulations with empirical data, we measured for each colony the proportion of simulations departing as far or farther from the colony average as did the experimental value. Twice this proportion gave the p-value for a test of the null hypothesis that the observed value was drawn from the same probability distribution as the simulated values.

#### Results

The results show no significant difference between experiment and simulation for five of six colonies (Table 4). This outcome validates our model’s ability to reproduce observed patterns of splitting in two-nest emigrations for a variety of colony compositions.

**Table 4:** Percentage of brood in the better nest for each of the six colonies, predicted vs observed. The last column is the p-value, with P <0.05 indicating a significant difference between predicted and observed percentages.

| Colony | % Pred | %Observed | P |
| --- | --- | --- | --- |
| A4 | 59 ± 17 | 61 | 0.86 |
| A6 | 61 ± 28 | 80 | 0.56 |
| A8 | 63 ± 30 | 99 | 0.36 |
| A14 | 59 ± 20 | 98 | 0.1 |
| A16 | 61 ± 34 | 100 | 0.5 |
| A17 | 60 ± 25 | 2 | 0.02 |

### 4.3. Two Unequal Nests: Speed-Accuracy Trade-off

The third question we ask is: does our model reflect the way that colonies trade off speed and accuracy when the urgency of the emigration changes? In “urgent” situations, ants face a critical need for immediate new shelter and thus benefit from moving out of the old nest location as fast as possible. In contrast, in less urgent situations they can deliberate longer among alternatives to increase the likelihood of moving directly to the best site (Pratt and Sumpter, 2006; Dornhaus et al., 2004). We simulated experiments that adjusted urgency by offering colonies a choice between a mediocre and a good nest under two circumstances: their old nest has just been destroyed (high urgency) or their old nest is of acceptable quality but worse than either of the new candidates (low urgency). Our simulations followed the same tactic by tuning the physical quality of the home nest to adjust urgency. We ran 300 simulations each for eleven home nest qualities in range [0,1], with candidate nest qualities of [1,2]. We used the default parameter values, except **lambda_sigmoid**, which was set to 16 in order to increase the ants’ sensitivity to home nest quality differences, and **pop _coeff**, which was set to 0 in order to better match the model assumptions in (Pratt and Sumpter, 2006). As in (Pratt and Sumpter, 2006), we measured the duration of emigration as the time in rounds at which the old site was completely abandoned, and we measured the accuracy of decision-making as the proportion of the colony’s members inside the good site at the time of old nest abandonment.

#### Results

The results show that time taken to complete an emigration decreased as urgency increased (i.e., as old nest quality decreased) (Fig. 5b). This is consistent with the empirical observation that higher urgency induces faster emigrations (Fig. 5a). Furthermore, the simulations show that higher urgency (lower old nest quality) reduces the likelihood of the colony achieving consensus on the better site. This also matches the empirical results, which show that higher urgency leads to lower accuracy (Pratt and Sumpter, 2006).

**Figure 5:**
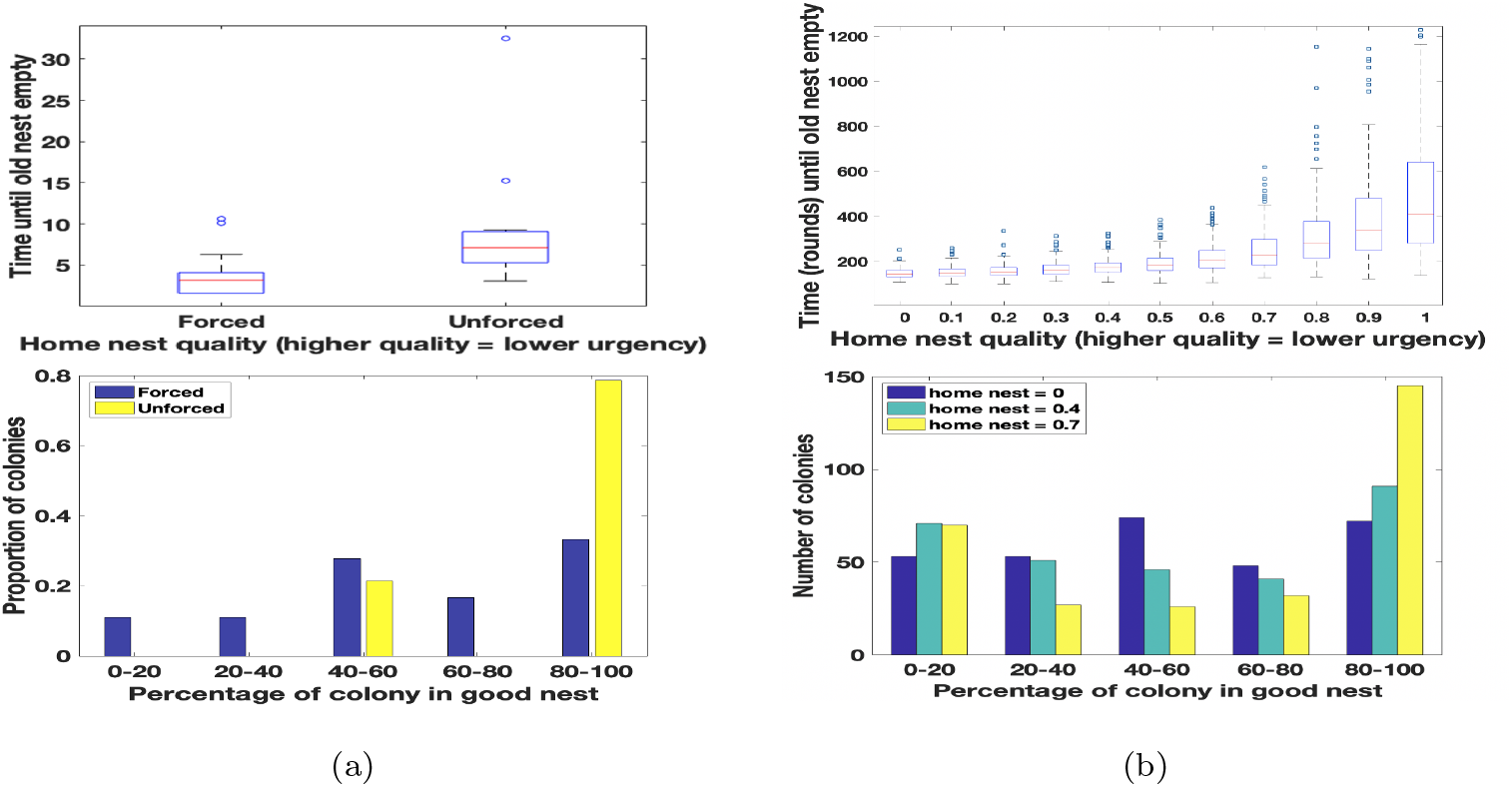
(a): From (Pratt and Sumpter, 2006): Speed and accuracy of decision-making in high-urgency and low-urgency emigrations. (Top) Time until the old nest is empty for each treatment. The ends of each box mark the upper and lower percentiles, and the horizontal line inside the box gives the median. The brackets show the data range, and circles are outliers. (Bottom) Histograms of the degree to which colonies split between the good and mediocre new nest sites, under high urgency (blue bars) and low urgency (yellow bars). (b): Simulation results for high and low urgency emigrations. Top and bottom panels correspond to those in (a). Simulations were run for more urgency levels than in the experiments.

These results confirm that our model can account for the empirically observed speed and accuracy trade-off up to old nest abandonment. However, it is worth noting that real colonies in the low urgency situation were better able to reach consensus than our simulated colonies. This might suggest the existence of other mechanisms at work that this simulation failed to capture. One such mechanism could be sensitivity to the presence of nestmates when assessing the quality of a nest. This could enhance consensus by amplifying the differential treatment of competing nests, as the better one’s population increases and makes it still more attractive. Our model can capture this phenomenon by using a non-zero *pop_coeff*, a possibility explored further in the next section.

## 5. Confirmation of New Experiments

In this section we consider more complex scenarios where the link between colony patterns and individual behavior has not previously been modeled.

For scenarios that have been explored empirically, we determine how well our model can account for observed results. Section 5.1 examines a colony’s ability to choose well when faced with larger option arrays; and Section 5.2 focuses on how colonies make rational decision time investments depending on nest quality differences.

### 5.1. Colonies Have High Cognitive Capacity

How well do colonies perform when selecting from many nests? A previous study (Sasaki and Pratt, 2012) showed that colonies are quite good at selecting a single good nest from a set of eight nests, four of which are good and four of which are mediocre. This is in contrast to individual ants, who are as likely to choose a mediocre as a good nest when faced with the same scenario. The colony advantage has been hypothesized to result from sharing the burden of nest assessment: very few of the scouts ever visit more than one or two nests, whereas a lone ant visits several, potentially overwhelming her ability to process information about them successfully. We simulate this experiment to determine whether we can reproduce both the colony’s ability and the observed distribution of nest visits across scouts.

We designed a simulated experiment with multiple nests in the environment, half of which are mediocre (physical_quality 1.0) and the rest of which are good (physical_quality 2.0). We considered three environments with 2, 8, and 14 nests, respectively. For each environment, We ran 600 simulations with a fixed colony size 200, containing 50 active and passive ants each, and 100 brood items.

#### Results

First, we found that simulated colonies reached consensus on a good nest with high probability, matching that seen in empirical data (Fig. 6). This was true even when the number of nests was increased to 14.

**Figure 6:**
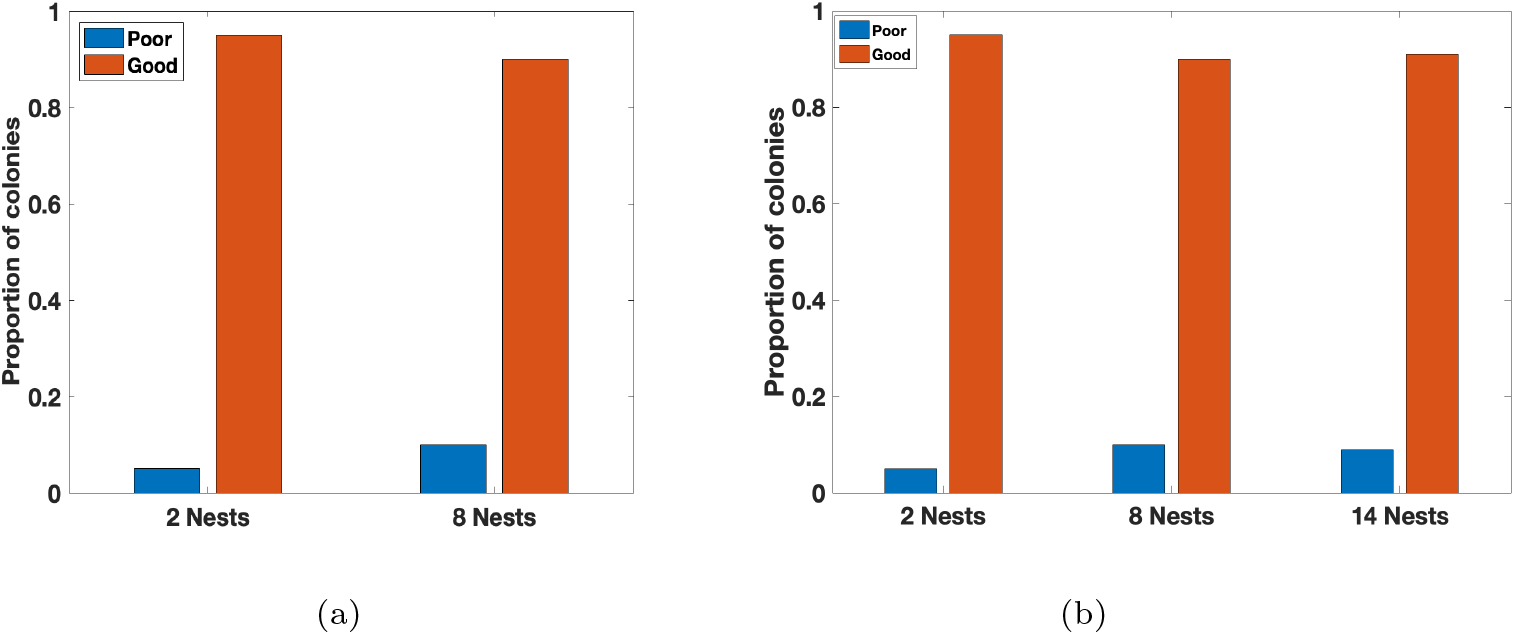
The proportions of colonies that eventually moved into poor or good nests. (a): Empirical results in 2-nest and 8-nest settings (Sasaki and Pratt, 2012). (b): Simulation results from our model in 2-nest, 8-nest, and 14 nest settings.

Next, we verified that the high cognitive capacity of colonies is associated with a low number of nests visited by each scout. The proportion of ants visiting only one or two nest was similar in the simulations and experiments (Sasaki and Pratt, 2012): over 80% of individual ants visited only one or two nests in the course of the emigration. Fig. 7 shows similar pattern is seen for the number of transports: that is, if we focus only on the ants who contributed to the emigration by transporting nestmates, over 80% visited only one or two nests. Thus, ants that access many nests have a minor role in the transportation process, supporting the hypothesis that colonies’ high cognitive capacity results from avoiding the overloading of individual ants.

**Figure 7:**
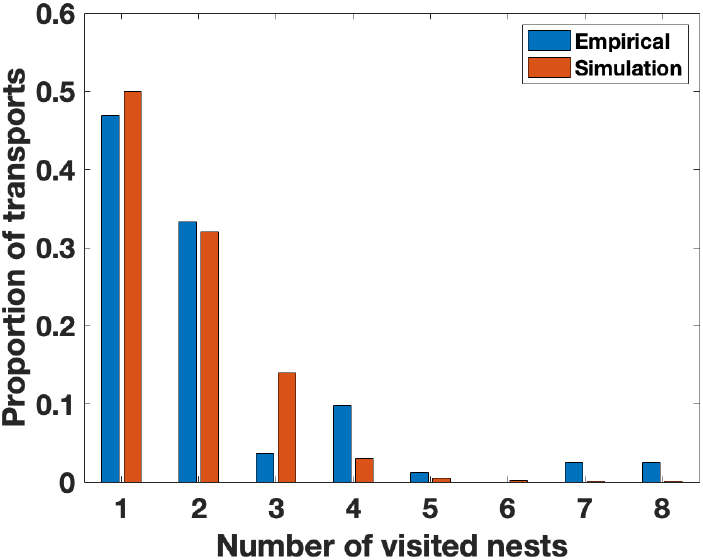
Proportions of transport efforts as a function of the number of candidate sites visited by each ant. The blue bars show the percentage of transports done by ants that visited a given number of nests (Sasaki and Pratt, 2012), and the dark orange bars show the same for simulated ants. Colonies choose among eight nests (four good and four mediocre) in both simulations and experiments (Sasaki and Pratt, 2012).

### 5.2. Colonies Make Rational Choices about Decision Speed

For choices between two nests, how does the difference between the nests affect the speed of decision-making? Counter-intuitively, a previous study (Sasaki et al., 2019) found that colonies move more quickly when site qualities are more similar. But this behavior accords with decision theory predictions that decision-makers should take less time if the consequences of their choice are small; that is, since the nests are similar in quality, the opportunity cost of making a wrong decision is small, so it’s rational to save time costs by taking on a higher risk of choosing the wrong nest.

We simulate this scenario to determine if we can reproduce the same pattern, but we also explore a broader range of quality differences to better describe the relation between quality difference and decision time. We designed an environment with two candidate nests, one good and the other mediocre. The good nest has physical_quality 2 in all simulations, but the physical_quality of the mediocre nest varies across simulations from 0.2 to 1.7. We asked whether the quality of the mediocre nest is correlated with the convergence score (a measure of decision speed). We ran 300 simulations for each environment with a colony of size 200, consisting of 50 active workers, 50 passive workers, and 100 brood items. We repeated this set of simulations for five different values of **lambda_sigmoid** values: [8,10,12,14,16].

#### Results

If our model reproduces the rational time investment choices of colonies (Sasaki et al., 2019), then we expect the convergence score to increase as the mediocre nest quality increases, thus becoming more similar to the good nest. Our results partially match this prediction, with convergence score increasing as the mediocre nest quality goes from 0.2 to about 1 (Fig. 8). However, at higher mediocre nest qualities, the pattern reverses and convergence score declines. This basic pattern is seen for all tested values of **lambda_sigmoid**.

**Figure 8:**
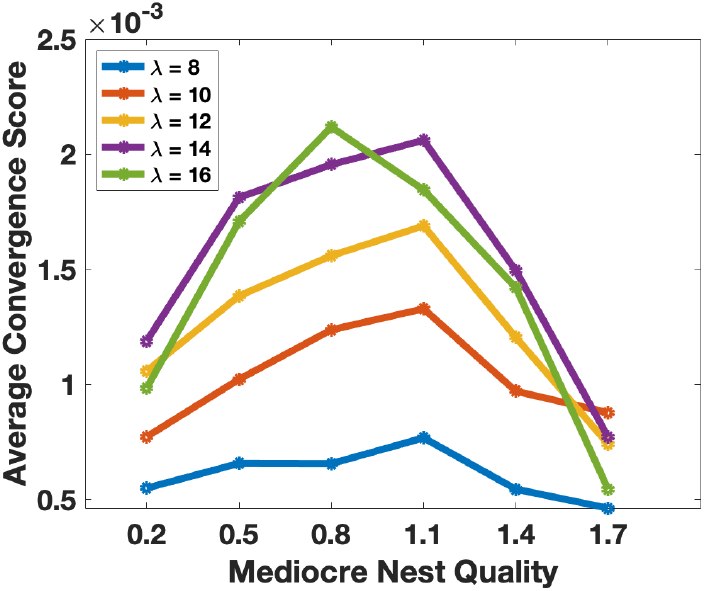
Average convergence score as a function of the physical-quality of the mediocre nest. The physical-quality of the good nest is 2, and that of the home nest is 0.

We propose that the nest qualities studied in (Sasaki et al., 2019) came from the region below the peak score that saw an increase of speed with decreasing quality difference. But from our more granular simulations, we predict that as the quality difference gets still smaller, the convergence score will start decreasing, meaning colonies will start investing more time.

Why might this happen? Recent studies have explained the behavioral difference between individuals and colonies via two different decision models: the tug-of-war model describes individual behavior, while colony behavior is better accounted for by the horse race model (Kacelnik et al., 2011). The tug-of-war correctly predicts the irrational behavior of individual ants, in that their decision-making slows down for options that are more similar. The horse race, in contrast, correctly predicts colonies’ rational acceleration of decision making for similar options. We hypothesize that the applicability of these models to the colony’s behavior changes as the quality difference changes. More specifically in Fig. 8, before the peak score is reached, the colony may effectively distribute its decision-making across many ants with limited information, the situation envisioned in the horse-race model. After the peak score is reached, the colony may come to depend more on individual comparisons between nest sites made by a few well-informed ants, and thus to show the irrational slow-down predicted by the tug-of-war model. It could also be the case that more transports are performed between the two candidate nests as the likelihood of the mediocre nest achieving quorum attainment increases.

## 6. The Power of Social Information

In this section, we explore the influence of social information on migration speed, accuracy, and cohesion. Section 6.1 explores correlations between **pop_coeff** and the degree of randomness in individual decision-making; and Section 6.2 reveals how **pop_coeff** decreases splitting by colonies facing two equal options.

### 6.1. Balancing Personal and Social Information

Individual ants are capable of directly comparing nests and choosing the better one, but their discriminatory ability is less than that of whole colonies. This may be seen as a kind of “wisdom of crowds,” in which the estimations of many noisy individuals are integrated into a more precise group perception. Ants do this via positive feedback loops based on recruitment, which can amplify small differences in site quality (Sasaki et al., 2013). They also use social information via the quorum rule, under which full commitment to a site is conditioned on a minimum number of nestmates “voting” for it by spending time there. The quorum rule inspired us to consider another way that ants might use social information to improve decision-making: by taking population into account when assessing a site’s quality. We do this via the parameter **pop_coeff.** which controls the degree to which the presence of nestmates increases a site’s perceived value. We propose that this population sensitivity might be able to complement the noisy perception of individual ants, modeled by the parameter λ in the Eq. 2. We hypothesize that ants may adapt to different values of **lambda_sigmoid** by changing the value of **pop_coeff.** In particular, we sought evidence for a correlation between the values of **lambda_sigmoid** and **pop_coeff** needed to achieve the best convergence score.

To investigate this question, we ran simulations for different combinations of **pop_coeff**(ranging from 0.002 to 0.8) and **lambda_sigmoid**(ranging from 2 to 16). We ran simulations for an environment containing two identical new nests [0,1,1]. For each combination of **pop_coeff** and **lambda_sigmoid**, we ran 500 simulations with a colony of size 200, consisting of 50 active workers, 50 passive workers, and 100 brood items.

#### Results

The results show evidence for an inverse relation between **pop _coeff** and **lambda_sigmoid** (Fig. 9). For each value of **lambda_sigmoid** in the range [2,16], there is a value of **pop_coeff** that maximizes the convergence score, and this value increases as **lambda_sigmoid** decreases. Thus, when an individual ant makes noisy local decisions (modeled with lower values of **lambda_sigmoid**), she can counteract this deficiency by relying more on the input of her peers through a higher value of **pop_coeff**.

**Figure 9:**
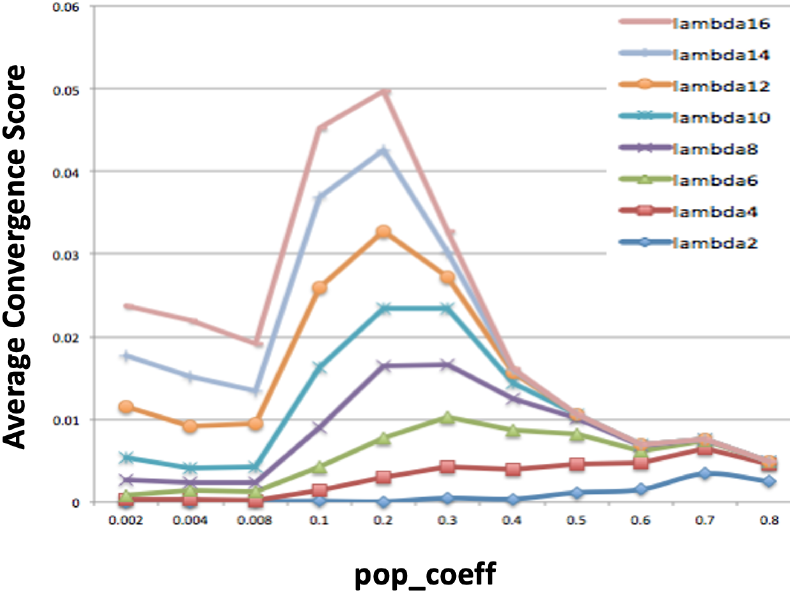
Average convergence score (across 500 simulations) as a function of **pop_coeff**. Different colored curves represent different **lambda_sigmoid** values as described in the individual decision model (Fig. 2). This shows that the optimal value of **pop_coeff** increases as **lambda_sigmoid** decreases.

However, a high value also has risks. Here we interpret the advantages and disadvantages of *increasing* the value of **pop _coeff**:

##### Advantages

- Higher momentum in the system. This can promote the colony to accumulate population at a certain nest more quickly, and thus achieve faster convergence.
- Better prevention of splits. Multiple candidate nests may reach the quorum, especially when the nests have similar physical qualities. This can lead to the colony splitting between more than one site. Social information via **pop_coeff** might help to break ties, by amplifying small random differences in the populations of competing sites.

##### Disadvantages

- Slower error correction. Since we are dealing with a randomized algorithm, there is always a chance that the colony will collectively make a “bad” temporary decision, even if individuals have low noise levels. The higher momentum will then make the wrong decision more “sticky” by accumulating more ants at a mediocre nest even if a better one is available. The colony would then have to move later to the better nest, adding costs in time and risk. In this way, high **pop_coeff** can cause slower convergence, and lead to “madness of the crowd”.

These trade-offs suggest that there is an optimal value of **pop_coeff** for a given **lambda_sigmoid** as seen in Fig. 9. This predicts that colonies may tune **pop_coeff** according to the uncertainty of individual behavior in order to achieve the highest convergence score for a given environment.

### 6.2. Avoiding Splits Between Two Equal Nests

In this section we further explore how social information can help colonies to reach consensus when faced with two identical nests. Many social insects have highly nonlinear recruitment mechanisms that lead to symmetry breaking when faced with two identical resources. For example, ant species that recruit via trail pheromones will choose one of two identical food sources rather than forming trails to both. This is because the attractiveness of a trail is a sigmoidal function of the amount of pheromone it contains, which leads to rapid amplification of small random differences in the strengths of competing trails (Beckers et al., 1990; Perna et al., 2012). However, similar experiments on *Temnothorax* ants found that they do not always break symmetry, instead exploiting both feeders equally, a result that has been attributed to the linear relationship between tandem running effort and recruitment success (Shaffer et al., 2013).

An open question is whether this lack of symmetry breaking also holds for nest site selection. When presented with identical nests, do colonies choose only one or split between them? If they can reach consensus, then how do they do so? One possibility is that the quorum rule provides sufficient non-linearity to amplify small random differences in site population, thus ensuring that the colony does not split. Another possibility is that colonies have some other as of now unrecognized mechanism of avoiding splits. A good candidate for such a mechanism is incorporation of site population into each scout’s assessment of site quality, as discussed in Section 6.1. This would allow amplification of early random differences in population, by increasing the likelihood of recruitment to the nest with more ants. We explore this question by simulating emigrations in which a colony is presented with two identical nest sites. We assess how well they reach consensus on a single one. We also vary the degree of scout sensitivity to site population by considering different values of **pop_coeff.**

We ran 200 simulations each for **pop_coeff** = [0, 0.1, 0.2, 0.3, 0.4], in an environment with **nest_qualities** = [0,2,2]. We set **lambda_sigmoid** to 16 in order to be more sensitive to temporal differences in nest populations. From an initial set of simulations, we observed that almost all simulations converged within the default value of **num_rounds**. Therefore, in order to gain more insight into the effect of **pop_coeff** on the degree of splitting before convergence, we set it to a smaller value 1000. The rest of the parameters take the default values.

#### Results

The simulation results show strong symmetry breaking (Fig. 10). That is, a large majority of simulations ended with 80% to 100% of the colony in one of the two nests. When consensus was reached, it was roughly equally likely to be in nest 1 or nest 2, producing the distinctive U-shaped distribution seen in Fig. 10. This pattern was true regardless of the value of **pop_coeff,.** suggesting that the quorum rule is enough to generate symmetry breaking in this case. However, as the value of **pop_coeff** increases, the histograms also aggregate more towards the two end bins, meaning there are fewer split cases. Thus we confirm the positive effect of **pop_coeff** in reducing splits, either by prevention or by facilitating later re-unification. These mechanisms can have significant effects in more challenging environments with more candidate nests.

**Figure 10:**
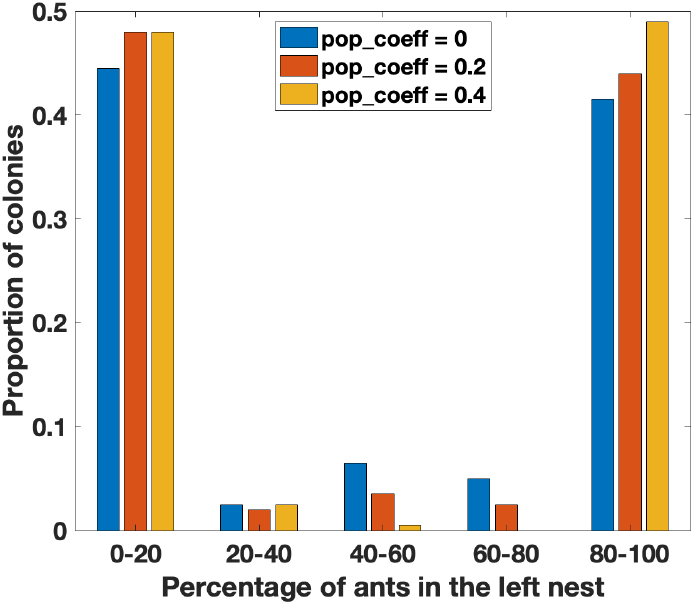
Simulation results for colonies choosing between two identical nests. The histograms show the distribution of the percentage of the colony occupying the left nest, for three different values of **pop_coeff**.

## 7. New Predictions

We also use our model to develop new hypotheses and predictions for future experimental study. Section 7.1 gives simulated evidence for a surprising speed-accuracy trade-off for the entire emigration process, tuned by the quorum size; and Section 7.2 discusses colony re-unification after splits with an increasing level of difficulty.

### 7.1. Quorum Size and the Speed/Accuracy Trade-off

*Temnothorax* colonies can adjust their behavior to adaptively trade off the speed and accuracy of decision-making (Pratt and Sumpter, 2006; Dornhaus et al., 2004) (Section 4.3). One of the behavioral tools implicated in this adjustment is the quorum rule. Colonies lower the quorum in more urgent situations, increasing their reliance on individual judgement. This allows them to make rapid decisions and quickly move the colony out of the old nest, at the cost of an increased probability of splitting or choosing an inferior nest (Franks et al., 2003a, 2009).

When considering speed, previous studies focused on the time to move out of the old nest, but the completion of an emigration often requires more than that. A fast “first” decision does not always mean a fast emigration. In fact, a low quorum and hence a fast “first” decision could lead to slower emigrations (Franks et al., 2009) since it could cause more splitting, which the colony must subsequently resolve in a second phase of movement. Here, we explore the effect of quorum size on the speed and accuracy as we have defined them for the whole process (Section 3.2). Within the accuracy measure, we pay special attention to the rate of splitting, which is the percentage of emigrations that do not reach a persistent convergence within the given number of rounds. A natural question arises: is there a speed-accuracy trade-off if we define “speed” as (the inverse of) the time taken to the final completion of the emigration? In other words, do the convergence score and accuracy have inverse correlations with **quorum_thre**, and are these relationships affected by splitting rate?

We simulated an environment with candidate nests [0.5, 1, 1.5, 2] and a home nest with quality 0 as usual. We used a colony of size 200, consisting of 50 active workers, 50 passive workers, and 100 brood items. Quorum size is assumed to be proportional to the total number of adults in the colony, and is set to **quorum_thre** × **num_adults**. We varied **quorum_thre** from 0.03 to 0.39, and set **pop_coeff** to either 0 or 0.35. We set *num_rounds* to 2000 and ran 100 simulations for each unique combination of **quorum_thre** and **pop_coeff.**

#### Results

The simulation results show that the convergence score generally has a reverse-U shape that peaks at **quorum_thre** = 0.24-0.27 (Fig. 11a, Fig. 11b). In addition, the accuracy measure has a similar shape, but peaks roughly at **quorum_thre** = 0.1-0.15. The split rate, in contrast, has a U-shape, with a trough around **quorum_thre** = 0.15 to 0.18 (Fig. 11c and 11d).

**Figure 11:**
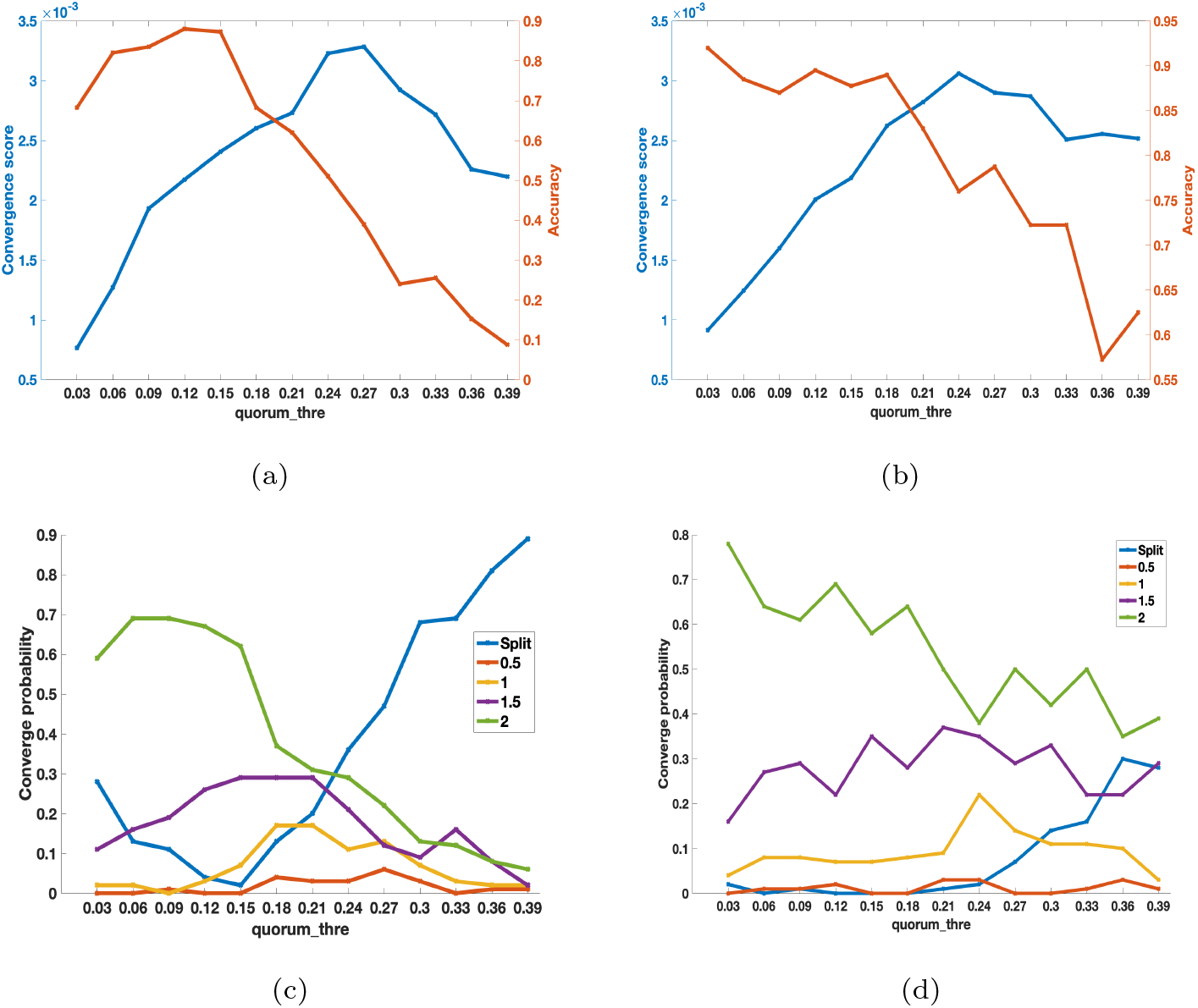
(a), (b): Convergence score and Accuracy as a function of **quorum_thre**, with **popcoeff** = 0 and 0.35 respectively. (c), (d): Probabilities of converging to each nest (or splitting) as a function of **quorum_thre**, with **popcoeff** = 0 and 0.35 respectively.

The above results indicate a surprising speed-accuracy trade-off in the segments where the two lines form an “X” shape in Fig. 11(a) and (b): the increase of **quorum_thre** is accompanied by a decrease in accuracy and an increase in speed. This is the opposite of our findings in Section 4.3 and in the related experimental work (Pratt and Sumpter, 2006; Dornhaus et al., 2004). However, it is important to note that the current definitions of speed and accuracy differ from those used in the prior work, which defined both quantities only up to the point where the old nest is empty. The results on splitting rate could give more insight into the conflicting results - if repairing splits is costly, lowering the probability of splits by increasing the quorum would indeed significantly increase the average convergence score. But another factor is that setting the quorum too high to reach will also delay convergence. These results point to the need for better understanding of how colonies reunite after splits, as well as the costs of reunification relative to other components of the emigration.

### 7.2. Reunification after Splitting

Finally, we touch on another aspect of the robustness of the house hunting algorithm — reunification after splitting. Experimental studies on the speed-accuracy trade-off showed that colonies often split in urgent emigrations, but they also noted that split colonies were eventually able to reunite (Franks et al., 2003a, 2009). Later studies (Doering and Pratt, 2019, 2016; Doering et al., 2020) showed that artificially divided colonies readily re-unite, using the same behavioral tools as in emigrations, but relying more on the efforts of a small group of active workers. These findings suggest that emigrations depend on a mixture of individual and colony-level decision making.

In this section, we explore how well our model achieves convergence after an arbitrary division among multiple nests. What can we learn about the mechanisms that achieve re-unification?

We ran simulations in which colonies were randomly divided among 2 to 9 nests. At the start of a simulation, each ant’s *location* variable in her *ExternalState* was sampled uniformly at random from all *env-choices*. We ran one set of simulations in which one nest was of quality 2 and the rest were of quality 1, and another set in which one nest was of quality 1 and the rest were of quality 2. We ran 300 simulations for each environment with a colony of size 200, consisting of 50 active workers, 50 passive workers, and 100 brood items.

#### Results

As the number of equal quality nests increases, the reunification task becomes increasingly difficult. Additional candidate nests have a negative effect on the convergence score and accuracy of reunification even when they are significantly worse than the best nest in the environment, possibly due to more distractions during evaluations of all nests. But the marginal effect of each additional nest diminishes (Fig. 12). As a result, the convergence score eventually stabilizes.

**Figure 12:**
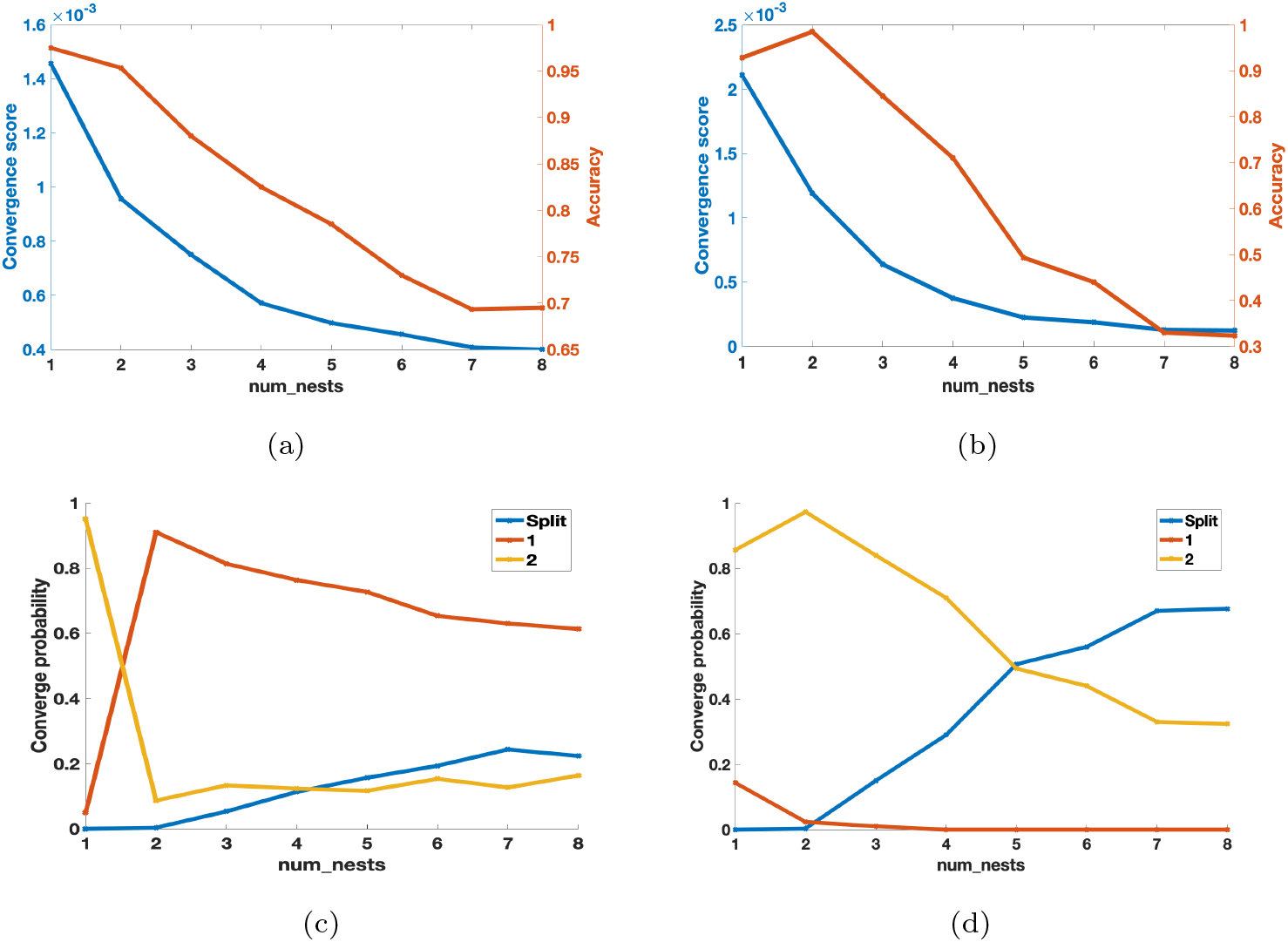
Convergence scores and splitting for environments with different numbers and qualities of nests. (a) Convergence score and accuracy as a function of the number of nests with quality 1 in the environment. (b) Convergence score and accuracy as a function of the number of nests with quality 2 in the environment. (c), (d) Same environments as (a) and (b), respectively, but plotting convergence probabilities to different nests (or splitting) on the y-axis.

However, we see that adding nests of quality 2 (highest quality in the environment) makes reunification much harder since split rate increases quickly. Intuitively speaking, having multiple nests that are the highest quality nest in the environment can greatly intensify competition among them. But this hypothesis needs additional quantitative analyses and empirical confirmation.

In these simulations we randomized the location at the start of our simulations, but not other variables in the internal and external states of individual ants. In reality, when ants are distributed among multiple nests, they most likely have a variety of values for these other variables. We further hypothesize that 1) randomizing the other variables may help with reunification, and/or 2) colonies may have mechanisms to prevent splitting to this extent during the emigration. However, further investigation is needed to test these hypotheses.

## 8. Discussion

In this paper, we introduce an agent-based modeling framework that can be used to formally represent a variety of distributed algorithms used by animal groups to understand the emergence of collective intelligence. We used the framework to examine the collective nest site selection process in colonies of *Temnothorax* ants. To test against existing experimental data and to make predictions, we built a convenient Python simulator that can easily be extended to add extra features. Our model reveals several underlying mechanisms behind the flexible, noise-tolerant and efficient collective house-hunting process in ant colonies. These include the mixed applicability of the horse-race model and the tug-of-war model to the house-hunting case; the colony’s emergent ability to tolerate noise and prevent splits by balancing social information and personal information; and the impact of splitting on the speed-accuracy trade-off of the emigration. Moreover, the generalizable modeling framework that we present can be used to investigate many other collective behaviors in biological systems.

The model successfully replicated published colony-level outcomes, suggesting that it accurately accounts for underlying individual behavior. It performed as well as an earlier model (Pratt et al., 2005), but did so with a more concise and elegant set of individual decision rules. The new model was also able to replicate more realistic and challenging emigration contexts that have not previously been modeled. Thus, on top of matching previous studies of simple one- and two- choice environments, our model successfully and quantitatively predicts colony behavior in environments with many more choices. Additionally, with varying degrees of experimental support, our model makes speculations on the quantitative relationships between multiple parameters and their effects on the colony’s collective behavior. The most important highlight is a novel role of social information in the speed, accuracy, and the rate of splitting during emigrations. The model and the accompanying software are versatile and easy to extend to additional investigations on unexplored scenarios such as emigrations in a changing environment. However, modeled colonies proved relatively weak at resolving splits in more challenging environments (i.e., with the colony split among several similar high-quality sites). This may reflect real limitations of the ants’ algorithm, but it seems more likely that the model underestimates their reunification abilities. This topic has so far received little attention; further development of this model could help to guide new experimental work.

In the rest of this section, we discuss several specific directions for future research. While our model captures many aspects of individual behavior, it leaves out some important features, including many that affect timing. These include 1) effects of the spatial distribution of nests, 2) effects of individual experience on recruitment probability and speed, and 3) actions that may last a variable duration such as the evaluation of a new nest. Adding these to the model would allow it to explore a broader range of colony abilities and to reveal as yet unknown components of individual behavior and how they interact with known aspects. For example, more realistic models of timing would undoubtedly affect the discovery behavior currently captured by the single variable *search_find*.

On the model analysis side, there are many directions for further research. First, we note the link between the effects of different quorum sizes and the horse-race and tug-of-war models that have been successfully used to describe group and individual decision-making, respectively, in these ants (Kacelnik et al., 2011). Our model finds that group decision-making may be better captured by the tug-of-war model when a colony is choosing between two very similar options. If so, this suggests that colonies can change their relative reliance on individual decision-making according to the decision context. This indicates the value of developing a more quantitative model that combines the tug-of-war and the horse-race models, based on the same factors that affect how a colony chooses the most beneficial quorum size.

Additionally, our model shows the potential utility of individual ants taking account of site population when assessing a site’s quality. Whether real ants use social information in this way has not yet been experimentally tested. Our results suggest that it may be important for preventing and repairing split decisions. However, the amount of social information that individuals should rely on is an intricate balance, as we described in Section 6.1. It would be highly valuable to quantify the relationship between the frequency and degree of splitting to the quorum size and to **pop_coeff**. A related research direction is to find out other factors that allow colonies to robustly reunify in split cases. However, the runtime of our program over hundreds of simulations can be significant, making it difficult to investigate the system dynamics and performance in all possible parameter settings. Overcoming this challenge requires software optimization techniques such as code parallelization and possibly further model simplifications.

On the theoretical analysis side, our model serves as a stepping stone for more rigorous mathematical formulations and proofs of guaranteed bounds on any metrics of interest. Starting with simpler environments, our model can be reduced to analytically derive the goals different mechanisms can and cannot achieve. These results can then potentially provide insights on why certain collective behaviors have emerged through evolution, as well as on engineering artificial distributed systems subject to similar limitations to reach consensus.

Finally, we emphasize that our modeling framework can be flexibly adapted to other distributed algorithms inspired by animal groups. One compelling example is that honeybee colonies use a very similar algorithm in their nest-selection process (Laomettachit et al., 2016), and can be easily modeled by our framework. Comparing it to our ant colony model can reveal commonalities and differences in how different animal groups achieve various goals and organize potentially conflicting priorities.

## 9. Acknowledgements

We thank Anna Dornhaus and Frederik Mallmann-Trenn for discussing the background literature and possible research directions early in the project. We also thank Emily Y. Zhang and Lili Su for reviewing the manuscript and providing valuable feedback.

## 10. Supplemental Materials

### 10.1. State Transition Details

#### Exploration

- An ant in *at_nest_E_* has four possible actions. First, she can perform “no_action” and remain in the current nest. Second, she can perform “search” and go into the state *search_E_*. Third, she can receive a “lead” by another ant to follow a FTR to a destination nest, *ec* ∈ **env-choices**, in which case she sets *old_candidate_nest* to the value of *candidate_nest*, and sets *candidate_nest* to *ec*. Then she transitions to the state *follow_E_*. Finally, she can receive a “carry” by another active worker ant to a destination nest *ec* ∈ **env-choices**, in which case her *location* and *candidate_nest* are changed to *ec*, and she stays in *at_nest_E_*.
- An ant is in the state *follow_E_* if she is in the middle of following an FTR, and has two possible actions. First, she can successfully follow the FTR to the destination nest (“follow_find”) and change her *location* to her *candidate_nest*, which results in the state *arrive_E_*. Otherwise, she may lose contact with her tandem leader (“get_lost”), and then enters the state *search_E_* and assigns the value of *old_candidate_nest* to *candidate_nest*.
- An ant in the state *search_E_* has three possible actions. First, she can have “no_action” and transition to *at_nest_E_* by staying at her last known *location*. Second, she can “find” a new nest, *ec* ∈ **env-choices**, in this round, assign the value of *candidate_nest* to *old_candidate_nest* and assign *ec* to both *location* and *candidate_nest*, and transition into *arrive_E_* state to evaluate it further. Third, she can receive an action, “carry”, and the results are the same as receiving the “carry” action in *at_nest_E_*.
- An ant in the state *arrive_E_* has two action options. First, she can “reject” the nest she just arrived at. She then assigns the value of *candidate_nest* to *location* and then that of *old_candidate_nest* to *candidate_nest* go into the *search_E_* state. Otherwise, if she performs “no_reject”, she transitions into the state *at_nest_A_* and assigns the value of *candidate_nest* to *location*.

#### Assessment

- An ant in the state *at_nest_A_* is assessing a new nest and is currently located at that nest. From here, three actions are available. First, she can “accept” the nest if she deems it high quality, which results in her transitioning to *at_nest_C_*. Second, she may perform “search” to get into the *search_A_* state. Third, she can receive a “lead” by another ant to follow a FTR to a destination nest, in which case she assigns the value of *candidate_nest* to *old_candidate_nest* and assigns the destination nest *ec* ∈ **env-choices** to *candidate_nest*, and then she transitions to the state *follow_A_*. Finally, she can receive a “carry” by another active worker ant to a destination nest *ec* ∈ **env-choices**, in which case her *location* and *candidate_nest* are changed to *ec* and transitions back to *at_nest_E_*.
- An ant in the states *follow_A_* or *search_A_* has the same options and variable changes as in *follow_E_* or *search_E_* respectively, but the resulting state sub-scripted with *A* except the “carry” action.
- An ant in *arriveA* state has the same options and variable changes as in *arrive_E_*, but with “reject” action leading to *search_C_*.

#### Canvassing

- An ant in *at_nest_C_* state has three available actions. First, she can decide to “recruit” and go into *quorum_sensing_C_* state. Second, she can decide to “search” more and result in *search_C_* state. Third, she may receive a “carry” by another active worker ant to a destination nest, in which case her *location* and *candidate_nest* are changed to that nest and results back to *at_nest_E_*.
- An ant in *quorum_sensing_C_* state is at a nest different than her home nest, and has two options. If she estimates the current nest population to be higher than the quorum threshold, she performs “quorum_met”, swap the values of *home_nest* and *candidate_nest*, and enters the state *transport_T_*. Otherwise, she performs “quorum_not_met” and enters *lead_forward_C_* state.
- An ant in *lead_forward_C_* state has three actions available to her. First, she can “lead” another active worker and lead her on an FTR from the original home nest to the candidate new nest. She changes her *location* to the value of *candidate_nest*, and enters *at_nest_C_* state. Second, she can “get_lost” in the process if she loses contact with the follower, and enters *search_C_* state. Lastly, she can “terminate” her emigration if the termination conditions are met, namely if she has repeated attempts to call other active workers who are also in *lead-forward_C_* state. In this case, she changes her *location* to her *home_nest*, resets *terminate_count* to 0, and enters state *at_nest_E_*.
- An ant in *search_C_* state has the same options and variable changes as in *search_E_* with the resulting state sub-scripted with *C*.
- An ant in *arrive_C_* state has the same options and variable changes as in *arrive_E_*, but with “reject” action leading to *search_C_*.

#### Transport

- An ant in *at_nest_T_* state has the same options and variable changes as in *at_nest_C_* with the resulting state sub-scripted with *T*, except that a “recruit” action results in *transporte_T_*, and that it can receive one additional action “lead”, in which case she assigns the value of *candidate_nest* to *old_candidate_nest*, assigns the destination nest *ec* ∈ **env-choices** to *candidate_nest*, and transitions to *follow_T_*.
- An ant in *transport_T_* state has three available actions. First, she can decide to carry another ant, active, passive, or brood, to her newly committed nest. This results in her entering *reverse_lead_T_* mode, meaning she can lead a reverse tandem run (RTR). These are tandem runs lead from the newly committed nest to the old home nest or another nest. Second, she can decide to “stop_trans” and stops her transport to go into the state *search_T_*. Third, similar to the state *lead_forward_C_*, there is a “terminate” action when the termination condition is met, namely if she has repeated attempts to carry other active workers who are also in *transport_T_* state. In this case, she changes her *location* to her *home_nest*, resets *terminate_count* to 0, and enters state *at_nest_E_*.
- An ant in *reverse_lead_T_* only has two actions as her options. First, she may perform *no_action* and returns to *at_nest_T_* state. Second, she may experience “delay” in her tandem runs, and will stay in *reverse_lead_T_* state. There’s no conclusion on the purpose of RTRs at this point in the research community, so we model it as a round-trip from an agent’s candidate nest to the original home nest and back, eventually ending up with no state changes.
- An ant in the states *follow_T_* or *search_T_* has the same options and variable changes as in *follow_E_* or *search_E_* respectively, but the resulting state are sub-scripted with *T* except the “carry” action.
- An ant in *arrive_T_* state has the same options and variable changes as in *arrive_A_*, but with “reject” action leading to *transport_T_*.

### 10.2. Simulation Details

#### 10.2.1. Sample Configuration File

~~~
[ENVIRONMENT]
num_ants = 200
nest_qualities = 0,1,2
~~~

~~~
[ALGO]
lambda_sigmoid = 8
pop_coeff = 0.35
quorum_thre = 0.15
quorum_offset = 0
search_find = 0.005
follow_find = 0.9
lead_forward = 0.6
transport = 0.7
~~~

~~~
[SETTINGS]
plot = 0
total_runs_per_setup = 500
num_rounds = 4000
percent_conv = 0.9
persist_rounds = 200
~~~

#### 10.2.2. Data Structures and Global Variables

We define four native data structures, as shown in Fig. 1. The global variables include 1) the transition tables defined in Fig. 3, 2) *Nests*, the array of all nests including the home nest which by default has quality 0 and id 0, and 3) *Ants*, the array of all ants in the colony.

#### 10.2.3. Simulation Overview

We describe our algorithm implementation in details below. Our executable software and instructions are available upon request.

Consider a colony of size *num_ants* where all the ants start the house-hunting task synchronously. We divide the total time to completion into *rounds*, with a maximum round number of *total_runs_per_setup*. At the beginning of round *t*, no ant has transitioned yet (instantiate 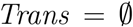). Then a random permutation of all *ant_ids* is generated as the order of execution. When an ant gets her turn during this round, she first checks if her *ant_id* is in *Trans*. If so, she does nothing. Otherwise, knowing its id and current state, she chooses an action for this round according to the probability distribution defined in the **select-action** function.

The action picked by an ant *x* has an *action_type*, a receiving ant id, and a nest id. Please note here that in real ant colonies, an action can involve either just a single ant, or a pair of ants (tandem run and carry). In the single ant action case, the receiving ant’s id is assigned value −1. In the pair ant action case, the action includes the valid *ant_id* of the receiving ant *y*. Similarly, not all actions require a nest, in which case the nest id for the action is −1.

By looking up the *Ants* array, *x* can also get the current external state of all ants including the receiving ant *y*, if any, of the picked action. These values are enough for *x* to call the **transition** function, and adds its own id to *Trans*. The special case handling is detailed in Section 2.2.3, including the case where *y* might also call a **transition** function and adds itself to *Trans*.

When one round finishes, each ant has had one chance to initiate or receive an action, and potentially has a new state. Repeat rounds like the above until the criteria is met for convergence with persistence, or until the program reaches the maximum number of rounds specified in the configuration file.

